# Learning the high-dimensional immunogenomic features that predict public and private antibody repertoires

**DOI:** 10.1101/127902

**Authors:** Victor Greiff, Cédric R. Weber, Johannes Palme, Ulrich Bodenhofer, Enkelejda Miho, Ulrike Menzel, Sai T. Reddy

**Affiliations:** Department of Biosystems Science and Engineering, ETH Zürich, Basel, Switzerland; Institute of Bioinformatics, Johannes Kepler University, Linz, Austria; Health & Environment Department, Molecular Diagnostics, AIT – Austrian Institute of Technology, Vienna, Austria

## Abstract

Recent studies have revealed that immune repertoires contain a substantial fraction of public clones, which are defined as antibody or T-cell receptor (TCR) clonal sequences shared across individuals. As of yet, it has remained unclear whether public clones possess predictable sequence features that separate them from private clones, which are believed to be generated largely stochastically. This knowledge gap represents a lack of insight into the shaping of immune repertoire diversity. Leveraging a machine learning approach capable of capturing the high-dimensional compositional information of each clonal sequence (defined by the complementarity determining region 3, CDR3), we detected predictive public- and private-clone-specific immunogenomic differences concentrated in the CDR3’s N1-D-N2 region, which allowed the prediction of public and private status with 80% accuracy in both humans and mice. Our results unexpectedly demonstrate that not only public but also private clones possess predictable high-dimensional immunogenomic features. Our support vector machine model could be trained effectively on large published datasets (3 million clonal sequences) and was sufficiently robust for public clone prediction across studies prepared with different library preparation and high-throughput sequencing protocols. In summary, we have uncovered the existence of high-dimensional immunogenomic rules that shape immune repertoire diversity in a predictable fashion. Our approach may pave the way towards the construction of a comprehensive atlas of public clones in immune repertoires, which may have applications in rational vaccine design and immunotherapeutics.

## Introduction

The clonal identity, specificity, and diversity of adaptive immune receptors is largely defined by the sequence of complementarity determining region 3 (CDR3) of variable heavy (V_H_) and variable beta (V_β_) chains of antibodies and TCRs, respectively [1–5]. The CDR3 encompasses the junction region of recombined V-, D-, J-gene segments as well as non-templated nucleotide (n, p) addition [6]. Due to the enormous theoretical diversity of antibody and TCR repertoires (>10^13^) [7–10] and technological limitations (Sanger sequencing), it was long believed that clonal repertoires were to an overwhelming extent private to each individual [11,12]. However, due to recent advances in high-throughput immune repertoire sequencing, it has been observed that a considerable fraction (>1%) of CDR3s are shared across individuals [1,5,13–26]. Thus these shared clones (hereafter referred to as “public clones”) are refining our view of adaptive immune repertoire diversity. Therefore, a fundamental question emerges: are there immunogenomic differences that predetermine whether a clone becomes part of the public or private immune repertoire?

In the context of antibody and TCR repertoires, the large theoretical clonal (CDR3) diversity renders the investigation of public and private repertoires computationally challenging [27]. Previous studies using conventional low-dimensional analysis suggested that public clones are germline-like clones with few insertions, thereby having higher occurrence probabilities, whereas private clones contain more stochastic elements (i.e. N1, N2 insertions) [17,23]. In order to investigate the composition of large numbers of sequences with the appropriate dimensionality, sequence kernels are increasingly used [28,29]. Sequence kernels are high-dimensional functions which measure the similarity of pairs of sequences, for example, by comparing the occurrence of specific subsequences (k-mers) in a high-dimensional space [30,31]. Supervised machine learning (e.g., support vector machine analysis) is an approach, which takes low and high-dimensional feature functions as input to find a classification rule that discriminates between two (or more) given classes on a single-clone level (e.g., public vs. private clones) [32]. In contrast to using conventional low-dimensional features to analyze immune repertoires, the coupling of high-dimensional sequence kernels to support vector machine (SVM) analysis may lead to greater insight into the immunogenomic structure of repertoire diversity; specifically the difference between public and private repertoires. As opposed to previous approaches [33], a key advantage of sequence-kernel based SVM analysis is the prediction-profile-based identification of CDR3 subregions that are most predictive for a respective class (public or private class) [30,31]. This approach may lead to predictive immunological and mechanistic insight into the immunogenomic elements that shape repertoire diversity.

In order to identify the immunogenomic differences between public and private antibody repertoires (Figure 1), we applied support vector machine analysis (Figure 1B) to six large-scale immune repertoire (antibody and TCR) sequencing datasets from mice and humans (Figure 1A). When using low-dimensional features (germline gene and amino acid usage, CDR3 subregion length) as the input for SVM analysis, prediction accuracy of private and public status reached maximally 66%, which only slightly improves on a random classifier (50%). However, when implementing a high-dimensional sequence-kernel (sequence composition) based support vector machine analysis, we were able to detect strong immunogenomic differences concentrated in the N1-D-N2 region in public and private clones, with a high prediction accuracy (balanced accuracy≈79–83%, Figure 1C). Our results unexpectedly signify that both public *and* private antibody repertoires contain predictive high-dimensional features that enable their accurate classification. Our SVM approach was sufficiently robust to be applied across repertoire studies with different library preparations and high-throughput sequencing protocols demonstrating their widespread applicability.

**Figure 1.**
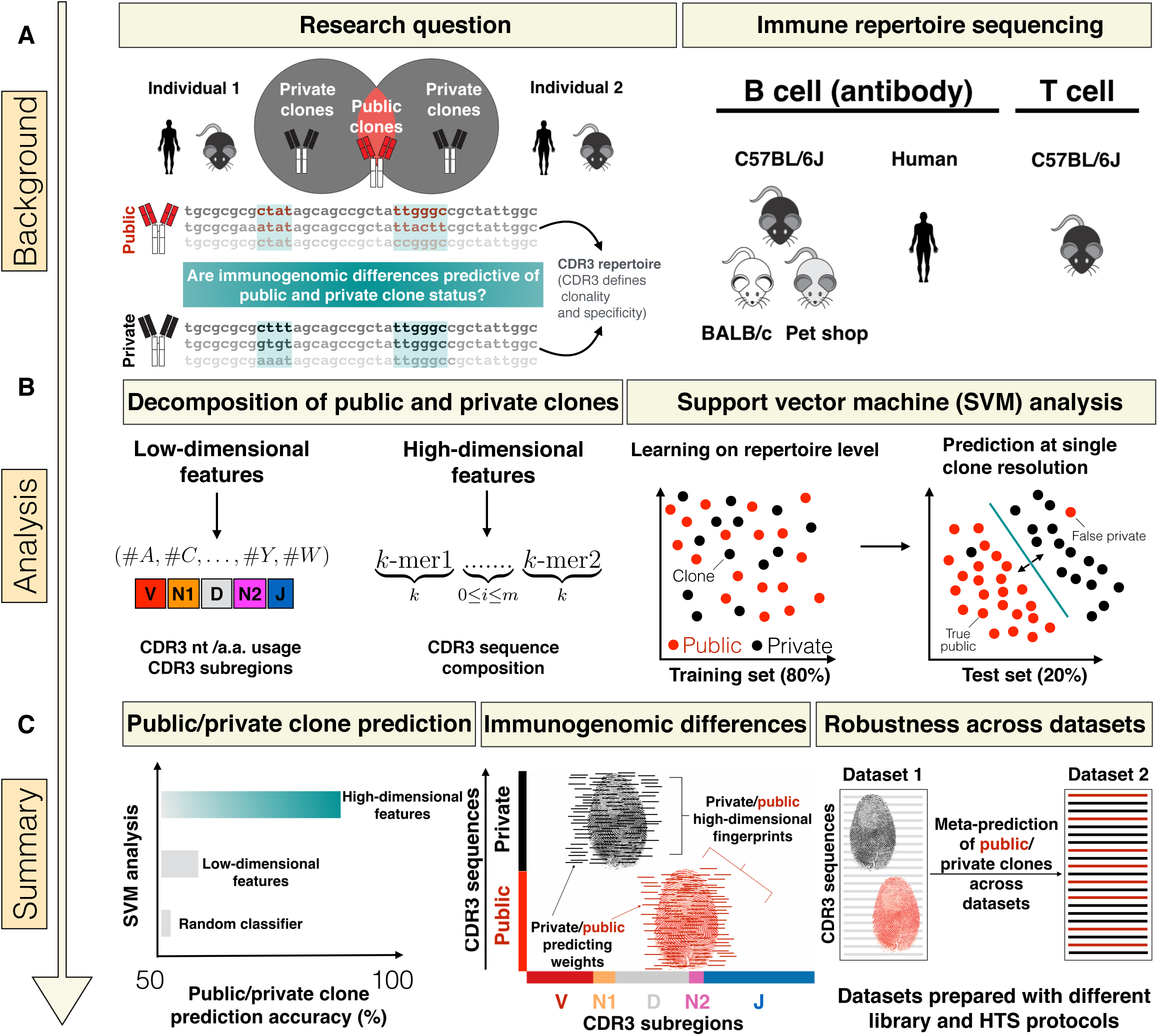
Immunogenomic analysis of public and private antibody repertoires. **(A)** We asked whether there are immunogenomic differences that predetermine a clonal sequence’s (CDR3) public or private status within a an immune repertoire. The public repertoire is composed of clones being shared among at least two individuals (we also explored an alternative public clone definition, Figure 6F). Private clones are those distinct to each individual. We defined antibody and T cell clones based on 100% CDR3 (complementarity determining region 3) identity. For statistical power, we used six large-scale datasets (Supplementary Table 1) comprising different B-cell populations, species (humans, mice) and immune antigen receptors (B/T cell receptor). **(B)** To answer our question, we decomposed public and private immune repertoires in conventional low-dimensional features (e.g., CDR3 amino acid usage, Figures 2 and 3) or novel high-dimensional features (CDR3 sequence decomposition into subsequences of length *k* (*k*-mers) separated by a gap of length *m*, Figures 4 and 5). Leveraging supervised machine learning (support vector machines), we tested whether low and high-dimensional features can detect immunogenomic differences between public and private repertoires (see Methods) and consequently can be used for prediction of public and private status at single clone resolution. **(C)** We found that low-dimensional features are poor predictors of public and private clone status. In contrast, we detected strong predictive immunogenomic differences, concentrated in the N1-D-N2 CDR3 subregion, between public and private clones using high-dimensional features. Thus, public as well as private clones each possess a class-specific high-dimensional immunofingerprint composed of class-specific subsequences that enables their classification with high accuracy. Our SVM approach was found to be generalizable across datasets produced in different laboratories with different library preparation and high-throughput sequencing (HTS) protocols.

## Results

### Public and private clone repertoires cannot be predicted by germline gene or amino acid usage

As the basis for elucidating the immunogenomic differences between public and private clones, we used a recently published high-throughput sequencing antibody repertoire dataset [16] (Dataset 1, *Methods*). This dataset contains ∼200 million full-length antibody V_H_ sequences derived from 19 different mice, stratified into key stages of B-cell differentiation: pre-B cells (preBC, IgM), naïve B cells (nBC, IgM), and plasma cells (PC, IgG). This dataset thus provided the important advantages of both high sequencing and biological depth (preBC and nBC represent antigen-inexperienced cells, while PC are post-clonal selection and expansion due to antigen exposure). Public clones, precisely defined here as CDR3 sequences (100% amino acid identity) occurring in at least two mice, were found to compose on average 15% (preBC), 23% (nBC), and 26% (PC) of antibody repertoires across B-cell stages (Figure 2A). As previously reported, we found that public clones are both biased to higher frequencies and are enriched in sequences from natural antibodies (Supplementary Figure 10) [17,24,34]. Throughout B-cell development, public and private clones used nearly identical V, D, J, VJ and VDJ germline genes (overlap >95%), which were at nearly identical frequencies in preBC and nBC (Spearman r≈1) and at varied frequencies in PC (Spearman r>0.5–0.8) (Figure 2B). Thus, neither public nor private clones showed any preferential germline gene usage. On average as well as at each CDR3 sequence position, higher frequency amino acids occurred more often in public clones (e.g.: A, C, D), whereas lower frequency amino acids could be found at higher frequency in private clones (e.g.: H, I, K) (Figure 2C, Supplementary Figure 2B). This observation held true across all B-cell stages (r = 0.5–0.76; p<0.05, Supplementary Figure 1A). Repertoire-wide absolute differences in amino acid usage between private and public clones were slight (0.2–1.4 percentage points, Figure 2C). To test whether these repertoire-level differences were sufficient to predictively discriminate between public and private clones on a single clone level, we employed supervised support vector machine learning (SVM) analysis (*Methods*, Figure 1B). For all SVM analyses in this study, in order to minimize classification bias, a dataset was constructed for each repertoire, which consisted of all public clones and an equal number of private clones from the repertoire (Supplementary Table 1) such that both public and private clones had identical CDR3 length distributions. Subsequently, the dataset constructed for SVM analyses was divided into 80% training sequences and 20% test sequences (Figure 1B, *Methods*). We found that amino acid usage was a suboptimal predictor of clonal status with a prediction accuracy ≤ 65% (Figure 2D) where prediction accuracy is defined as the mean (balanced accuracy) of specificity and sensitivity (see *Methods*), as described previously [30,35].

**Figure 2.**
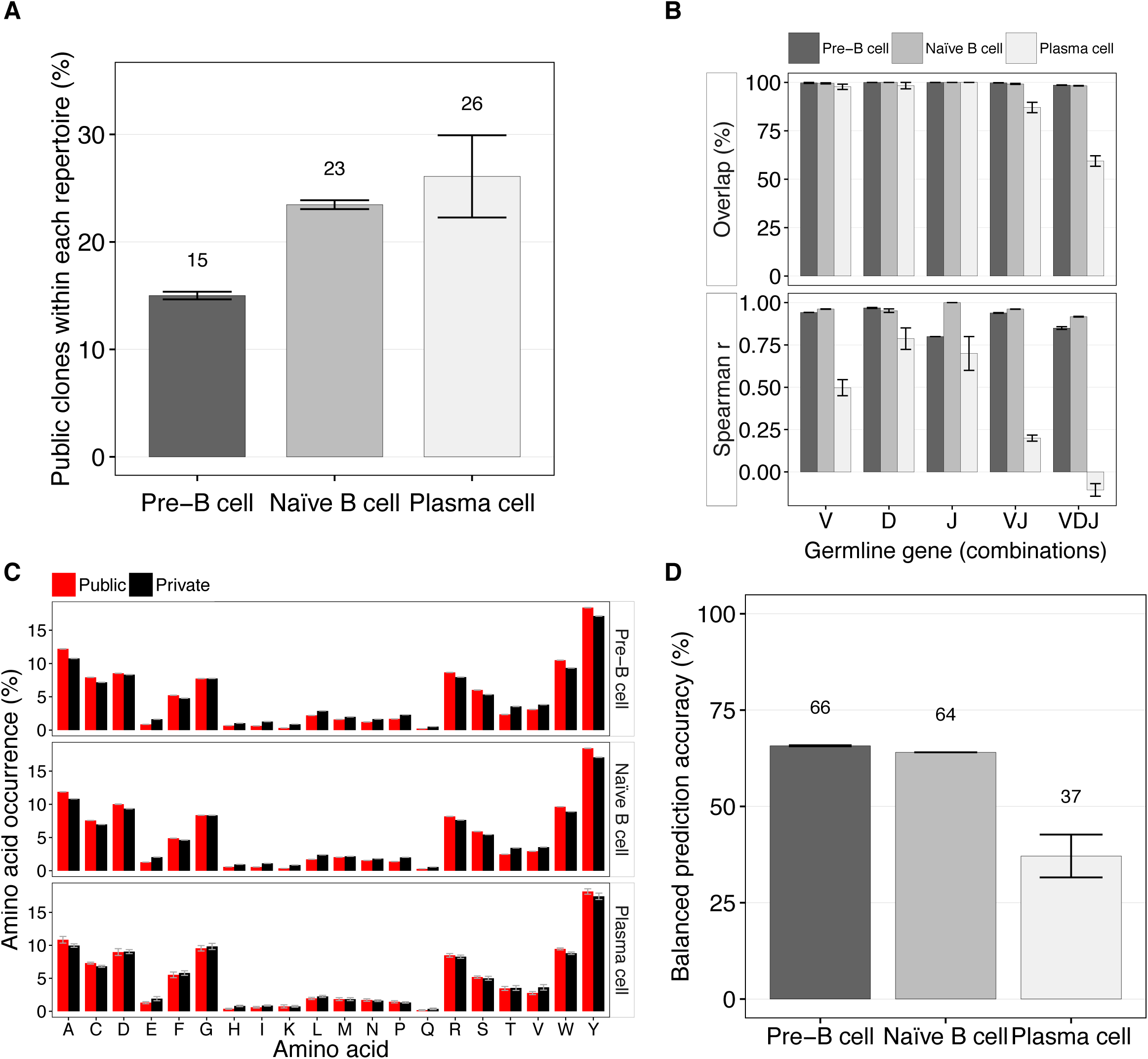
Public and private clone repertoires do neither differ predictively in germline gene usage nor amino acid composition. **(A)** Public clones represent 15–26% of murine antibody repertoires throughout B-cell ontogeny. Public clones were defined as being shared in at least two mice (see *Methods*). **(B)** Overlap and Spearman correlation of V, D, J germline genes and their respective combinations (V-J, and V-D-J) between private and public clones by B-cell population. **(C)** Relative amino acid composition of public (red) and private clones (black). Differences between public and private clones were not significant (Kolmorogov-Smirnov test, p>0.05). **(D)** SVM-based discrimination of public and private clones based on CDR3 amino acid composition (see Methods). Balanced prediction accuracy was defined as the mean of specificity (detection rate of public clones) and sensitivity (detection rate of private clones). Barplots show mean±s.e.m.

### Public and private clones do not differ predictively in CDR3 subregion length

Since public and private clones did not differ in germline gene usage, we asked whether they differed with respect to length and diversity of CDR3 subregions (V, N1, D, N2, J). The V, D and J subregions are derived from germline genes (IGHV, IGHD, IGHJ), while N1 and N2 represent the insertions introduced during the junctional recombination process (n- and p-nucleotides). Public clones in preBC and nBC repertoires possessed a relative V subregion length of 23–24% (Figure 3A), whereas private clones had slightly shorter V subregions (≈21%, p<0.05, Supplementary Figure 3A). The J subregion length behaved analogously (public: 40%, private: 36%) while the D subregion length did not differ between groups (public: 25%, private: 25%). We observed the largest difference between public and private clones in the relative length of N1 and N2 subregions with deviations of 36–46 percentage points from a 1:1 ratio (N1: public ≈6.5%, private ≈8.2%; N2: public ≈4.3%, private: ≈7.7%, p<0.05, Figure 3A, Supplementary Figures 3A, B). Conversely, PC CDR3 subregion lengths did not differ between public and private clones (with the exception of N1, which was slightly longer in public clones, Figure 3A, Supplementary Figure 3B).

**Figure 3.**
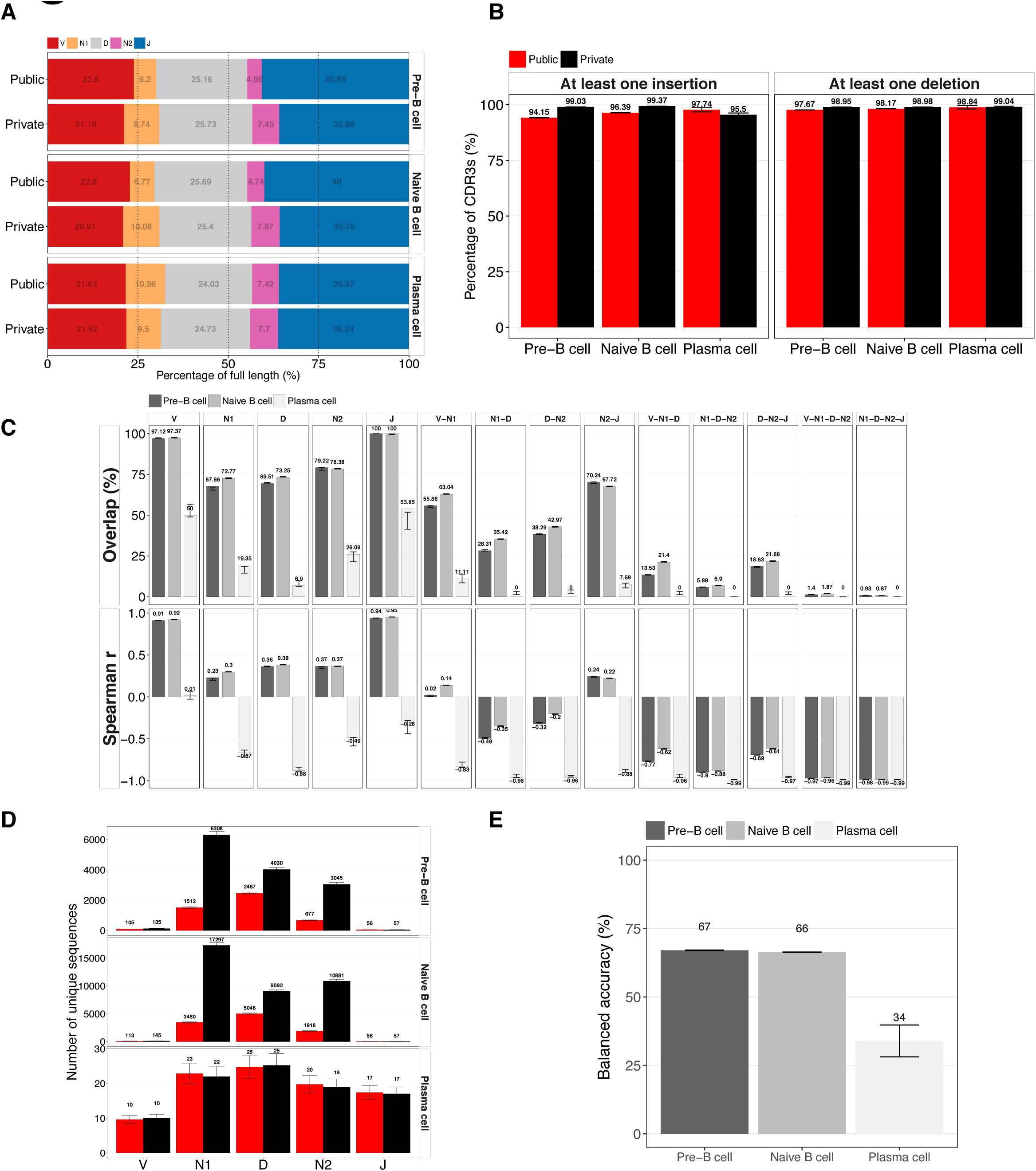
CDR3 subregion length does not predict a clone’s public/private status. **(A)** Normalized CDR3 subregion (V, N1, D, N2, J) lengths (median) of public and private clones by B-cell population. **(B)** Frequency of clones (public, private) with at least one N1/N2 insertion or deletion occurrence by B-cell population. **(C)** Overlap and Spearman correlation of CDR3 subregions and combinations thereof by B-cell population. **(D)** Number of unique V, N1, D, N2, J subregions (species richness) of public and private clones by B-cell population. Species richness of private clones CDR3 subregions was obtained by accounting for private and public clones size differences (bootstrapping, see *Methods*). **(E)** SVM-based prediction of public and private clones based on V, N1, D, N2, J subregion composition (Figure 3A, see *Methods*). Balanced (prediction) accuracy was defined as the mean of specificity (detection rate of public clones) and sensitivity (detection rate of private clones). Barplots show mean±s.e.m.

Regardless of public or private designation, nearly all CDR3s (>94%) had at least one nucleotide insertion (N1 or N2) and at least one deletion (Figure 3B), thus only a very small portion of clones were “germline-like” having neither insertion nor deletion (≤4%, Supplementary Figure 4C,D). Furthermore, across B-cell populations both N1 and N2 insertions were present in >50% and >70% of public and private clones, respectively. Of note, N1 and N2 insertions showed no preferential selection of germline gene segments (IGHV, D, J) (Supplementary Figure 3D) and the mean length of the sum of insertions (N1+N2) did not correlate with V-D-J frequencies (Supplementary Figure 5A, Pearson r=0).

Deletion length was highest in D subregions (mean of 5’ and 3’ D-deletions: ≈7 nt, Supplementary Figure 3C) whereas it was lowest in V subregions (≈0.8 nt, Supplementary Figure 3C). Although private clones showed a higher number of deletions, differences between public and private clones were slight (max difference≈0.6 nt, Supplementary Figure 3C). Of interest, we were unable to detect an association between the lengths of insertions and deletions (Supplementary Figure 5B).

Although differences in CDR3 subregion length and occurrence of insertions and deletions were significant in preBC and nBC (Figure 3A, Supplementary Figures 3A–C, 4C, D, p<0.05), training a SVM based on CDR3 subregion length, led to low prediction accuracy of public/private clone discrimination (balanced accuracy ≤ 68%, Figure 3E). This indicates that the slight differences observed in CDR3 subregion length on the repertoire level are not reliable for class prediction.

### Public and private clones show differences in sequence composition

Since *low-dimensional* features (CDR3 a.a. and subregion properties) did not achieve high discrimination accuracy between public and private clones (Figures 2D, 3E), we investigated whether CDR3 sequence composition (*potential dimensionality*: >10^13^ different CDR3 sequences) differed between public and private clones. In preBC and nBC, V and J subregions neither differed in public and private clones with regard to unique sequences (>97%) nor frequency thereof (Spearman r>0.95, Figure 3C). Consequently, we observed no differences in V and J subregion diversity (number of unique V and J subregions) between public and private clones (Figure 3D). Although there was a major difference in diversity of N1, D, and N2 subregions between private and public repertoires, as the number of private preBC and nBC clones surpassed that of public clones by 1.6–4.5-fold (Figure 3D, p<0.05, size adjusted, see also Supplementary Figure 3A), N1, D, N2 subregion overlap between public and private clones was >66% (Figure 3C). In PC repertoires, diversity differences between public and private repertoires were minimal but overlap of subregions reached maximally 46% and Spearman correlation was consistently negative. In contrast to single subregions, combinations of subregions showed low overlap between public and private repertoires irrespective of B-cell population (e.g., N1-D-N2 overlap in nBC was ≈6%, Figure 3C), which is explained by a large combinatorial diversity (Supplementary Figure 4B, Supplementary Table 2) of CDR3 subregions. Thus, sequence composition differed substantially between public and private clones.

### High-dimensional CDR3 sequence composition analysis predicts public and private clones with high accuracy

In order to test, whether the detected differences in sequence composition were predictive, we utilized high-dimensional sequence kernels for SVM analysis [30]. We used the gappy-pair sequence kernel [30,36,37], which decomposes each CDR3 into subsequences of length k (k-mers) separated by a gap of length *m* (Figure 4A, see *Methods*). Applying this kernel function to all CDR3s of a given training dataset generates a feature matrix of dimension *n*\**f*, which serves as input for the SVM analysis: here, *n* is the number of CDR3s in the training dataset and *f* the number of features. By cross-validation, we selected the parameter combinations that resulted in the highest prediction accuracy: *k*=3, *m*=1 at the nucleotide level (potential feature diversity: 8192, *Methods*) and *k*=1, *m*=1 at the amino acid level (potential feature diversity: 800, *Methods*). On both the nucleotide and the amino acid level, public and private clones in preBC and nBC could be classified with ≈80% accuracy, with very low variation across mice (Figure 4A, Supplementary Figure 6A, E). In order to validate the robustness of the chosen public clone definition, we showed that the SVM was *incapable* of separating public from public and private from private clones across individuals (balanced accuracy < 50%, Supplementary Figure 6D). In addition, we validated that the high prediction accuracy was maintained for an alternative and more stringent definition for public clones (balanced accuracy = 83–84%, Supplementary Figure 6F). In order to quantify the statistical significance of our high-dimensional SVM approach, we confirmed that the balanced accuracy was close to random (50%) when shuffling CDR3 nucleotide and amino acid sequences (Supplementary Figure 6B) and when shuffling public and private labels across clones (Supplementary Figure 6C).

**Figure 4.**
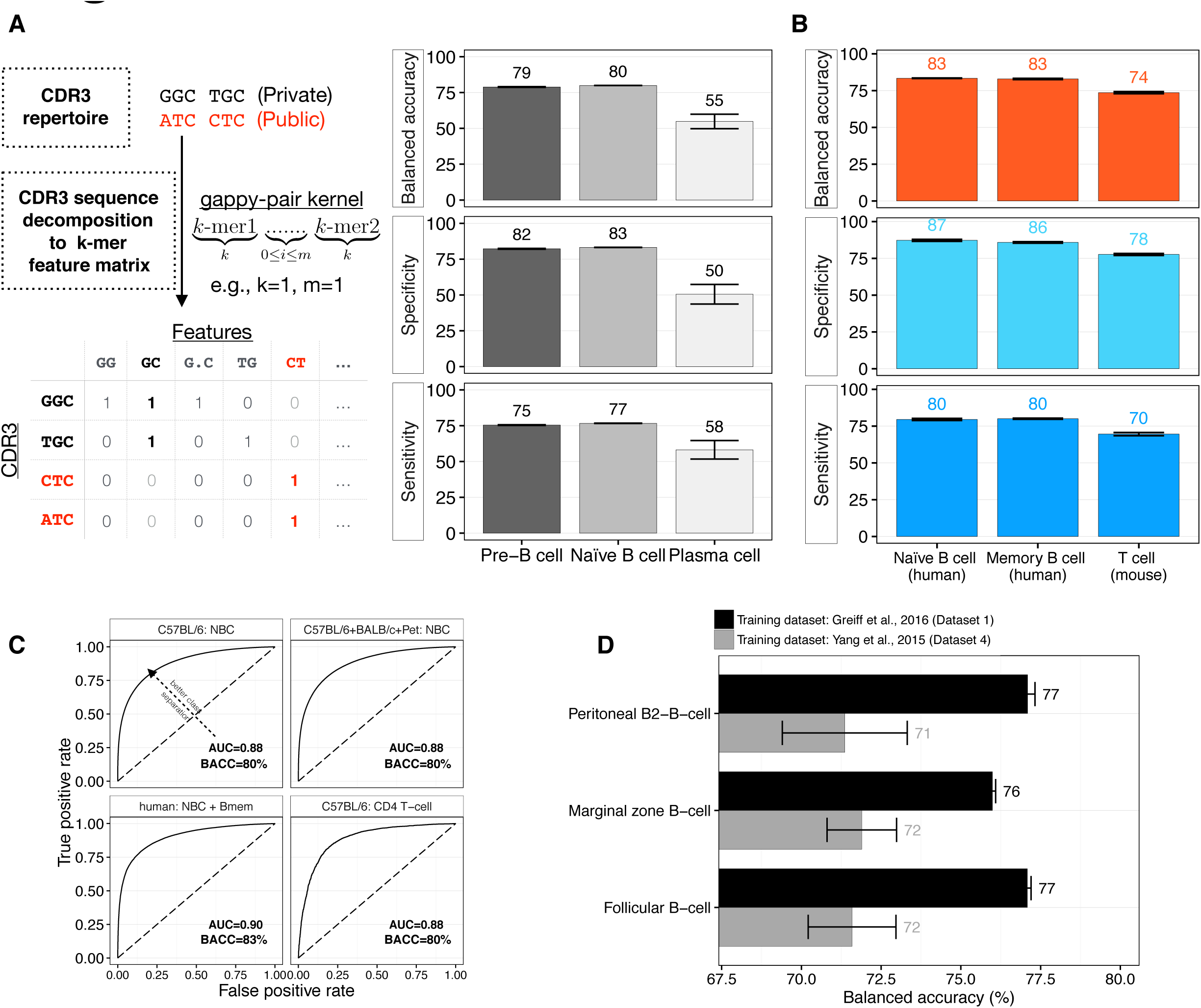
Public and private clones can be predicted with 80% accuracy using high-dimensional CDR3 sequence decomposition. **(A)** Specificity (detection rate of public clones), sensitivity (detection rate of private clones) and balanced accuracy (mean of specificity and sensitivity) for public vs. private clones SVM discrimination by B-cell population. For each repertoire, a dataset composed of equal numbers of public and private clones (nucleotide sequences, length equilibrated) was assembled (*Methods*, Supplementary Table 1). Subsequently, as displayed in the insert, the gappy pair kernel function decomposes each CDR3 sequence into features made of two *k*-mers separated by a gap of maximal length *m*. The maximal number of features is 4^(2×*k*)^×(*m*+1)=8192 for nucleotide sequences (*k*=3, *m*=1) and 20^(2×*k*)^×(*m*+1)=800 for amino acid sequences (*k*=1, *m*=1). Based on this decomposition, a feature matrix of dimension #CDR3s times #Features is constructed. Each row of the feature matrix thus corresponds to a feature vector for a CDR3 and contains counts of each feature as it occurs in the CDR3 sequence. These feature vectors serve as the input to the linear SVM analysis. The optimal parameter combinations (*k*=3/*m*=1 for nucleotide, *k*=1/*m*=1 for amino acid sequences) was determined by cross-validation on the training dataset (*Methods*). **(B)** Prediction accuracy of public vs. private clones of human naïve and memory B-cells, and murine T cells. SVM parameters were identical to those used in (A). **(C)** Public clones were accumulated across mice by B/T-cell populations (nBC, CD4), strain (nBC: C57BL/6, BALB/c, pet) or across B-cell populations (human naïve and memory B cells) in order to subsequently perform SVM-based classification as described in (A). Sizes of aggregated SVM-datasets ranged between ≈5x104 (CD4 T cell) and 3x106 (nBC: C57BL/6, BALB/c, pet) clones. ROC curves show excellent classification results (AUC [area under the ROC curve] ≈ 0.90). **(D)** SVM-based prediction of public vs. private clones across experimental studies. NBC repertoires of Dataset 1 (mean size: ≈180,000 clones) were used to predict public and private clones in the B2-B-cell repertoires of Dataset 4 (mean size: ≈2’400 clones, Supplementary Table 1). Barplots show mean±s.e.m.

Furthermore, we confirmed that the differences in immunogenomic composition between public and private clones were not exclusively mouse-strain-specific (C57BL/6); we replicated a balanced accuracy of ≈80% with repertoires from BALB/c and pet shop mice (Datasets 2, 3, Supplementary Figure 6A). Analogously, public and private clones could be discriminated with >80% accuracy in human B-cell repertoires (Figure 4B, Dataset 5). Finally, we showed that our approach also demonstrated high classification accuracy between public and private clones of mouse TCR Vβ repertoires (balanced accuracy = 74%, Figure 4B, Dataset 6).

Successful classification within each individual (mouse or human) proved that fundamental and stereotypical differences between public and private classes do indeed exist. However, theoretically, these differences could be specific to each individual and not generalizable. In order to exclude this possibility, we accumulated public and private clones across individuals into datasets of up to 3×10^6^ unique clonal sequences and showed that classification accuracy was maintained (Figure 4C), reaching a maximum in human naïve and memory B cells (balanced accuracy = 83%, AUC [area under the ROC curve] = 0.90). These results signified that the same set of features used to predict public and private clones within one individual is sufficient for prediction across individuals of the same species. Thus, the high-dimensional features provided by sequence kernels (800 for amino acid and 8192 for nucleotide) and learned on the repertoire level, were sufficient and generalizable to discriminate public from private clones in both humans and mice on a per clonal sequence basis (single clone resolution).

### Prediction by CDR3 sequence composition is dependent on dataset size and applicable across studies

Our high-dimensional sequence-composition-based SVM approach was unable to predict public and private clones in PCs (balanced accuracy = 50%, Figure 4A, Dataset 1). With respect to unique CDR3s, the PC SVM-dataset was 3 to 4 orders of magnitude smaller than that of preBC and nBC (Supplementary Table 1, Dataset 1); therefore we tested whether the low accuracy was due to sample size. We performed SVM analysis on datasets ranging in size from 100 to 230’000 unique CDR3 sequences (Supplementary Figure 7B) and found that prediction accuracy was indeed a function of sample size, increasing from 56% (100 clonal sequences) to 80% (230’000 clonal sequences). Thus, small sample size may explain the lower prediction accuracies observed in the PC (IgG) dataset. In further support of this hypothesis, we found that in a dataset of human memory B-cells (mixed IgM, IgG) (Dataset 5) that was 3 orders of magnitudes larger than the PC dataset, we were able to achieve >80% accuracy (Figure 4C), suggesting that prediction of public clones may also be possible for antigen experienced B-cell populations and is thus not limited to antigen-inexperienced ones.

Since we observed that dataset size was important for reaching high prediction accuracy (Supplementary Figure 7B), we asked whether cross-dataset meta-analysis, which leverages large datasets as training datasets for performing public and private clone prediction in other (smaller) datasets obtained from studies using slightly different library preparation and high-throughput sequencing protocols. To answer this question, we investigated the prediction accuracy of the sequence-composition-based SVM classifier trained on Dataset 1 (nBC B2-B-cell population), applied to a test dataset 100 times smaller (177’197 vs 1519 sequences), consisting of repertoires from various C57BL/6 B2-B-cell populations [20] (Dataset 4, Supplementary Table 1). By using the model based on the larger dataset (Dataset 1), prediction accuracy could be improved by up to 7 percentage points (76%–77% vs. 69%–73%, Figure 4D), which neared the prediction accuracy *within* Dataset 1 (Figure 4A). Thus, sequence-kernel-based SVM models can be effectively trained on large datasets (openly accessible) enabling robust predictive performance for meta-analysis *across* studies.

### Stereotypical immunogenomic differences between public and private clones are concentrated in the N1-D-N2 subregions

To identify the subregions that contributed most to classification accuracy, we performed sequence-kernel-based SVM on each CDR3 subregion separately as well as all ten relevant combinations thereof (Figure 5A). Classification based on each single or paired CDR3 subregions did not result in high prediction accuracy (balanced accuracy ≤ 67%, Figure 5A). Among the partial combinations, it was the N1-D-N2 subregion combination that achieved maximum prediction accuracy (74%, Figure 5A, Supplementary Figure 7A) approaching that of the full combination (V-N1-D-N2-J, ≈80%), indicating that the sequence composition between public and private clones differed most within N1-D-N2 subregions. J subregions contributed least to prediction accuracy as V-N1-D (balanced accuracy≈73%) and N1-D-N2 (balanced accuracy≈73%) surpassed D-N2-J (balanced accuracy≈70%, Figure 5A). In order to confirm that subregion differences between public and private clones were largely dictated by the N1, D and N2 subregions and not within the overhang regions linking N1, D, and N2, we showed that subregion shuffling impacted prediction accuracy only negligibly (Supplementary Figure 6E). Visually and numerically, we confirmed the N1, D, and N2 subregions to be the drivers of public and private clone discrimination by constructing prediction profiles, which quantify for each sequence the contribution of each position to the decision value (public, private). Differences in contribution to the decision value were highest in the sequence positions belonging to the N1, D, and N2 subregions (Figure 5B, Supplementary Figure 9). To summarize, our results indicate that the N1, D, N2 subregions of both public and private clone sequences contain class-specific predictive subsequences (k-mers) that enable the prediction of their status (public, private) with high accuracy.

**Figure 5.**
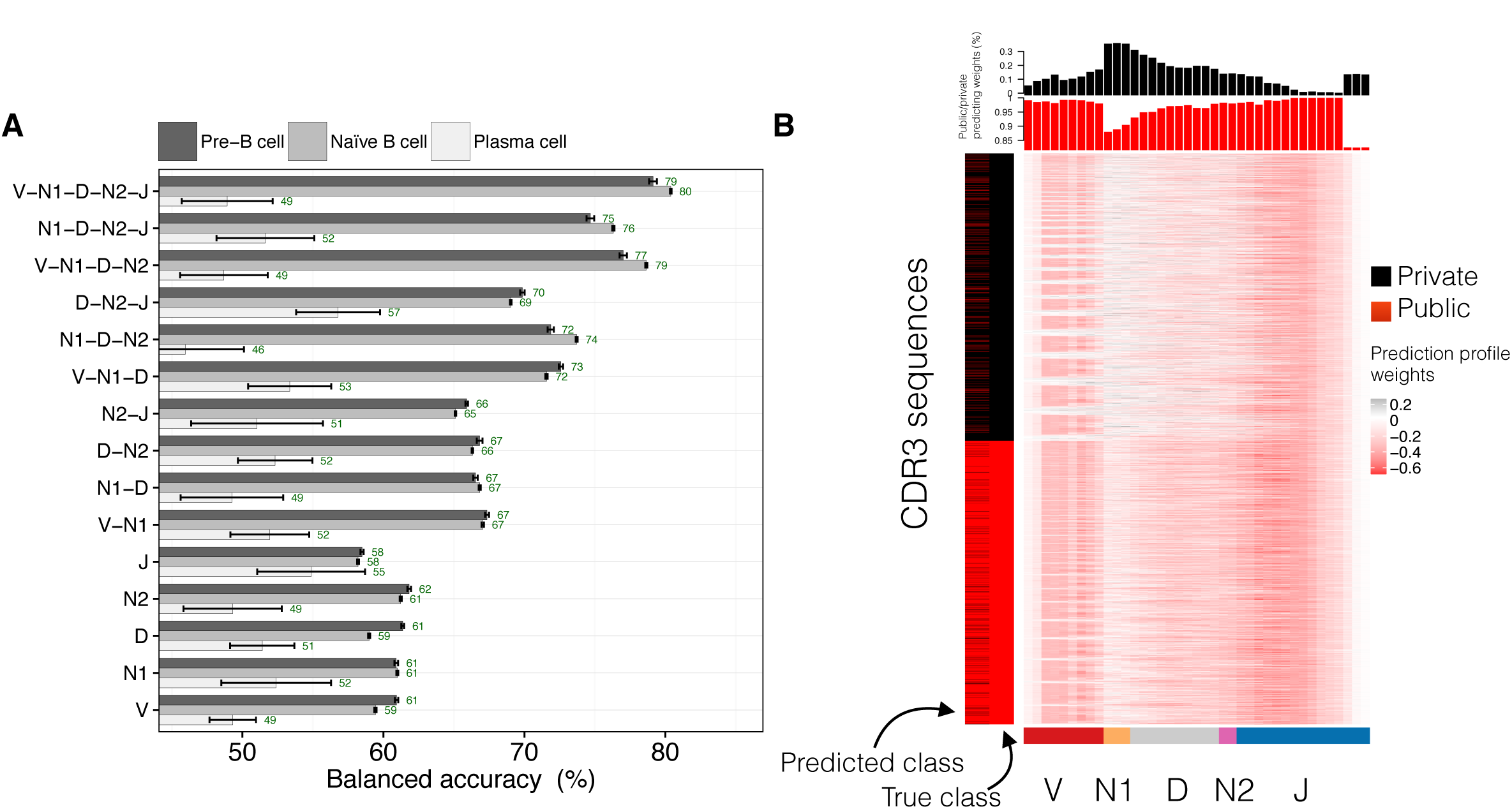
The N1-D-N2 subregions dominate the classification accuracy of public clones. **(A)** Balanced accuracy of public and private clone discrimination using sequence-kernel-basel SVM analysis. For each combination of CDR3 subregion, gappy pair kernel parameters (k, m, cost) were determined by cross-validation. Barplots show mean±s.e.m. **(B)** Exemplary visualization of prediction profiles of one test dataset (nBC) of CDR3s (rows) of length 39 (nt). Prediction profiles were computed as means of feature weights at each CDR3 position (1–39, see Methods). Positions colored red (<0) count towards “public” prediction of the respective CDR3s, whereas black-colored ones (>0) bias prediction towards the “private” clone status. Barplots indicate the percentage of private (black) or public predicting weights at each of the 39 positions. Color bars indicate median length of V (red), N1 (orange), D (grey), N2 (purple), J subregions (blue, Figure 3A). Prediction profiles across all CDR3 lengths as well as quantitative prediction profile analysis are given in Supplementary Figures 8 and 9, respectively.

## Discussion

We have performed a comprehensive immunogenomic decomposition of immune repertoires, which led us to conclude that *low-dimensional* features (Figures 2, 3, S1, S3–5) – CDR3 subregion length, germline gene usage, amino acid usage (Figures 2D, 3E) – were insufficient in detecting the immunogenomic shift between public and private clonal repertoires. In contrast, a *high-dimensional* sequence composition (sequence-kernel) approach could predict the public and private status of antibody clones within any individual with 80% accuracy. This CDR3 sequence-composition-based approach was generalizable across individuals, B-cell populations, mouse strains, species (mouse, human), immune cell types (B-cell, T-cell), and datasets produced in different laboratories (Figures 4B–D). While the appropriate definition of “public” clones is subject to current debate [5], the public clone definition adopted in this study has been used previously [5,22,38], and is the most lenient one possible. In fact, we showed that prediction accuracy only increased when increasing the stringency of the public clone definition (Supplementary Figure 6F). The fact that our SVM approach is robust to several public clone definitions, suggests there may not be the need for a consensus definition.

Sequence-kernel based machine learning analysis revealed stereotypical and predictive high-dimensional immunogenomic CDR3 subregion (N1, D, N2) composition biases (high-dimensional fingerprints) specific to both public and private clones, respectively (Figure 5). Those fingerprints achieved up to 100% prediction accuracy when isolated from V and J regions (Supplementary Figure 7A). Shuffling CDR3 subregions (V, N1, D, N2, J) impacted prediction accuracy only negligibly (Figure 5B, Supplementary Figures 6E, 9), confirming that N1, D, and N2 held the highest amount of class-specific information [25,39]. Of note, although the relative size of the human CDR3 N1-D-N2 subregion is larger than that of mice (≈65% [40] vs. 42% in mice, Figure 3A) with the N1-D-N2 subregion being the main amplifier of sequence diversity (Supplementary Table 2) [8,25], identical feature space sizes led to identical prediction accuracies for both species (Figure 4B). Thus, potential species-specific differences in sequence length and diversity did not impact the prediction accuracy of our approach. More generally, it is remarkable that feature spaces of dimension <10^4^ do not only suffice for detecting *sub*-repertoire clonal expansion-driven changes in individuals of different immunological status [29,35] but also provide ample combinatorial flexibility in defining fingerprints that discriminate *whole*-repertoire properties (public, private) within a >10^13^-dimensional space (Supplementary Table 2, [8,10]). This may point towards evolutionarily conserved traces in the immunogenome; for example, we found that murine public clones were enriched in natural antibody specificities (Supplementary Figure 10B).

Our results indicate that statistical significance does not necessarily translate into predictive performance: although CDR3 subregion length differed significantly between public and private clones (Figure 3A, Supplementary Figures 3A–C, 4C, D), the prediction accuracy of the low-dimensional SVM model based on CDR3 subregion length (Figure 3D) remained inferior to the high-dimensional one based on the actual sequence composition (Figure 4A). Furthermore, previous probabilistic work on modeling repertoire diversity indicated a broad range in clonal sequence generation probabilities – with (T cell) public clones suggested to be biased towards higher generation probabilities [24]. Corroborating these observations, we found that B cell public clones are more likely to have higher clonal abundance (Supplementary Figure 10A) – in general, however, public clones were distributed throughout the entire frequency spectrum from high to very low clonal frequency (Supplementary Figure 10A) [34]. Instead of attributing to each clonal sequence a generation probability, our work complements previous *probabilistic* work by leveraging a high-dimensional repertoire-level trained classifier for *binary classification* on a *per sequence basis*. It is this sequence-composition-based machine learning approach that led to the unexpected finding that also *private* clones – which were thought to be mostly stochastically generated – possess a high-dimensional fingerprint (predictive immunogenomic features).

Our SVM-driven approach enables rapid and accurate separation of large repertoire datasets into public and private repertoires. We note that mouse and human trained SVM-classifiers may not only be applied to experimental but also to synthetic repertoire data [41], which could pave the way towards the construction of a comprehensive atlas of human and mouse public clones. The high computational scalability of our machine learning approach – tested with as many as 3×10^6^ public and private sequences (Figure 4C) – allowed us to establish that the dataset size is a deciding factor for high prediction accuracy [33]: (i) in simulations, prediction accuracy increased by ≈30 percentage points when increasing the dataset size by 4 orders of magnitude from ≈10^1–2^ to ≈10^5^ clonal sequences (Supplementary Figure 7B). (ii) In experimental data, increasing training dataset size by 1–2 orders of magnitude (sequence data generated in a different lab using different experimental library preparation methods) increased prediction accuracy by up to 7 percentage points, suggesting large-scale cross-study detection of public clones is possible. (iii) The high prediction accuracy of human (antigen-selected) public and private memory B-cell clones (Figure 4B) suggested that the low accuracy of (antigen-selected) PC (IgG) repertoires (Figure 4A) may be due to small dataset size (Supplementary Table 1). More generally, we speculate that the prediction accuracies reported here merely represent lower bounds; future studies, which combine (i) advanced experimental and computational error correction methodologies (e.g., unique molecular identifiers) [42–44], (ii) high sampling and sequencing depth [1] and (iii) novel sequence-based deep learning approaches [45–47] may lead to even higher prediction accuracies.

To conclude, the existence of high-dimensional immunogenomic rules shaping immune repertoire diversity in a predictable fashion, leading to clones with higher occurrence probability within a population, highlights the potential of public clones to be a promising target for rational vaccine design and targeted immunotherapies [23,34,48,49].

## Methods

### Immune repertoire high-throughput sequencing datasets

We conducted our analysis on six high-throughput immune repertoire sequencing datasets, all of which are characterized below. Quality and read statistics can be found in the respective publications.

### Dataset 1

Murine B-cell origin (C57BL/6J): Sequencing data were generated by Greiff and colleagues [16]. B-cells were isolated from four C57BL/6 cohorts (n=4–5) including untreated and prime-boost immunized with protein antigens. Cells were sorted into the subsets pre-B cells (preBC), naïve B cell (nBC) and plasma cells (PC) by flow cytometry. Cell numbers per mouse were: 750’000 (preBC), 1’000’000 (nBC) and 90’000 (PC). RNA was isolated from cells, antibody libraries were prepared by RT-PCR and sequenced using Illumina MiSeq platform (2x300bp paired-end). The sequencing data has been deposited online (http://www.ebi.ac.uk/arrayexpress/experiments/E-MTAB-5349/) along with full experimental details and were preprocessed using MiXCR for VDJ-annotation, clonotype formation by CDR3 and error correction as described previously [16,50]. Briefly, for downstream analyses, functional clonotypes were only retained if: (i) they were composed of at least 4 amino acids, and (ii) had a minimal read count of 2 [51,52]. Public clones were defined as those clones that occurred in at least two different individuals within the same B-cell population and cohort.

### Dataset 2

Murine B-cell origin (BALB/c): Sequencing data were generated by Greiff and colleagues [16] and have been deposited online (http://www.ebi.ac.uk/arrayexpress/experiments/E-MTAB-5349/) with full experimental details. Briefly, naïve B-cells (1,000,000 cells per mouse) from 4 unimmunized BALB/c mice were isolated using the sorting panel from Dataset 1 and antibody libraries were prepared and sequenced analogously to Dataset 1. Data preprocessing was performed analogously to Dataset 1. Public clones were defined as those clones that occurred at least twice across mice.

### Dataset 3

Murine B-cell origin (Pet Shop mice): Sequencing data were generated by Greiff and colleagues [16] and have been deposited online (http://www.ebi.ac.uk/arrayexpress/experiments/E-MTAB-5349/) with full experimental details. Briefly, naïve B-cells (≈671’000 cells per mouse) from three pet shop mice were isolated and library preparation, sequencing, and data preprocessing was performed analogously to Dataset 1. Public clones were defined as those clones that occurred at least twice across mice.

### Dataset 4

Murine B-cell origin (C57BL/6J): Sequencing data were published by Yang and colleagues [20]: Mature B cells were extracted C57BL/6J-mice and sorted (1–2×10^4^ per cell population) into developmentally distinct subsets (splenic follicular B-cells (FOB, n=5), marginal zone B-cells (MZB, n=7), peritoneal B2-B-cells (n=5) and B-1a B-cells (n=43)). Data preprocessing was performed analogously to Dataset 1. Public clones were defined as those clones that occurred at least twice across mice of a given B-cell population.

### Dataset 5

Human B-cell origin: Sequencing data of naïve and memory B-cells from three healthy donors were published by DeWitt and colleagues [13] and downloaded already preprocessed from http://datadryad.org/resource/doi:10.5061/dryad.35ks2. Public clones were defined as those clones that occurred at least twice across individuals within a given B-cell population. Cell numbers of naïve and memory B cells were 2–4×10^7^ and 1.5–2×10^7^, respectively.

### Dataset 6

Murine T-cell origin: Sequencing data were published by Madi and colleagues [17]. CD4 T cells were isolated from 28 mice (three cohorts; untreated (n=12), immunized with complete Freud’s adjuvant (CFA, n=7) or immunized with CFA and ovalbumin (n=9). Data preprocessing was performed using MiXCR for annotation and error correction as described previously [16,50]. Public clones were defined as those clones that occurred at least twice across mice of a given cohort.

### Determination of statistical significance

Significance was tested using the Wilcoxon rank-sum test if not indicated otherwise. Where applicable, significance of correlation coefficients was tested using the R function cor.test() with default parameters.

### Statistical analysis and plots

Statistical analysis was performed using R [53] and Python [54]. Graphics were generated using the R packages ggplot2 [55], RColorBrewer [56], and Complex Heatmap [57]. Parallel computing of SVM analyses was performed using the R packages RBatchJobs [58] and doParallel [59].

### Definition of a clone

For all analyses, clones were defined by 100% amino acid sequence identity of CDR3 regions [1,16,51]. CDR3 regions were annotated and defined by MiXCR software [50] according to the nomenclature of the Immunogenetics database (IMGT) [60].

### Quantification of overlap

As defined previously [16], the percentage of clones shared between two repertoires *X* and *Y*: overlap 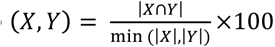, where |*X*| and |*Y*| are the clonal sizes (number of unique clones) of repertoires *X* and *Y*. A repertoire was mathematically defined as a set of unique clones.

### Junction Analysis

V, N1, D, N2 and J subregion annotation of sequences was performed using IMGT/HighV-Quest [61] (after initial preprocessing by MiXCR) [50]. Deletions were determined by finding the longest common substring between the germline genes and the V, D and J subregions identified in the CDR3 sequences.

### Estimation of the technological coverage of V, N1, D, N2, J regions

To estimate the technological coverage of each region (V, N1, D, N2, J), bootstrapping was conducted (Supplementary Figure 2). Briefly, 5, 25, 50, 75 and 100% of the full diversity of each region was sampled. Subsequently, the number of unique sequences per region present in the sample was compared to the total number of unique sequences.

### Determination of Shannon Evenness

The Shannon Evenness was calculated as previously described [35]. Briefly, we calculated the Hill-diversity for alpha = 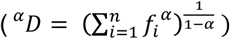 for a given frequency distribution (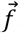, enumeration of the abundance of each subregion (combination)) of V, N1, D, N2, J subregions or combinations thereof. Subsequently, we obtained the Shannon Evenness ^*α*=1^=*E* by normalizing ^*α*=1^=*D* by the respective total number of V, N1, D, N2, J regions or combinations thereof (*n*) in the given repertoire.

### Estimation of the theoretical nucleotide diversity of the murine naïve clonal repertoire

The extent to which the entirety of the subregions V, N1, D, N2, J discovered in preBC and nBC of Dataset 1 covered any preBC/nBC repertoire was quantified by species accumulation curves as previously described [16]. Briefly, we defined the repertoire coverage (C_i_) of a given CDR3 subregion (R_i_) as the percentage overlap of its set of unique regions {R}_i_ with the set of regions contained in all previously accumulated repertoires 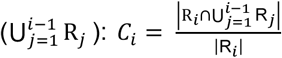, where *i* ∈ (1,…, *m*) with *m* being the total number of preBC and nBC repertoires (*m* = 38). To infer the number of subregions necessary for any given coverage, we used non-linear regression analysis using an exponential fit 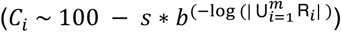 [62], where 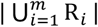 is the number of unique subregions contained within the accumulated repertoires and *s* and *b* are the parameters to be inferred. For ≥95% coverage, this is the estimated size of each murine naïve V, N1, D, N2, J subregion repertoire. We opted to report the coverage at 95% (Supplementary Table 2, column 2) to counter the effect of rare clones as described previously [16]. The product of the extrapolated coverage at 95% of each region (Supplementary Table 2) is the theoretical nucleotide diversity of the murine naïve clonal repertoire.

### Determination of private clones with high similarity to public clones

For each public clone, the number of private clones within 1 amino acid edit distance was enumerated (Figure 6B). Edit distance was determined using the stringdist() function (distance metric: Levenshtein distance) from the stringdist R package [63] as well as igraph [64]

### Support Vector Machine (SVM) analysis

In order to classify clones into public and private classes, a supervised learning approach was chosen in the form of a support vector machine (SVM) model. As input for all SVM analyses, CDR3-length equilibrated datasets were built for each sample (Supplementary Table 1). Briefly, for each sample, all public clones were paired in equal numbers with private clones of the same sample such that both public and private clones followed identical CDR3 length distributions. SVM analysis was performed using kernel-based analysis of biological sequences (KeBABS) [30] and sklearn [65], both of which are described in more detail below. For all SVM analyses, each dataset was split into training (80%) and test subset (20%). Cross-validation and SVM training was performed on the training dataset and class prediction on the test dataset. Prediction accuracy of class discrimination was quantified by calculating the balanced accuracy 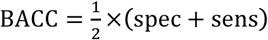, where specificity was defined as 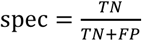, and sensitivity defined as 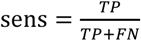 (*TP* = *True Positive, TN* = *True Negative, FP* = *False Positive, FN* = *False Negative*). Additionally, AUC (area under the curve, ROC curve) was calculated using the KeBABS R package [30]. An AUC value of 1 means perfect prediction accuracy (BACC = 100%), while an AUC value of 0.5 (BACC = 50%) is equivalent to random guessing.

### KeBABS support vector machine analysis

To discriminate public and private clones based on CDR3 sequence, we used the KeBABS R package [30], which implements kernel-based analysis of biological sequences. For all datasets, we used the position-independent gappy pair kernel [36,37], which divides all sequences into features of length *k* with gaps of maximal length *m* (Figure 4A). For the analysis of nucleotide sequences the parameters were set to *k*=3, *m*=1, C = 10, whereas the analysis of amino acid sequences was performed using parameters *k*=1, *m*=1, C = 100 (as determined by cross-validation). The cost parameter C sets the cost for the misclassification of a sequence. The maximal number of possible features used in the gappy kernel is determined by 4^2×*k*^×(*m* + 1) = 8′192 for nucleotide sequences and *20*^2×*k*^(*M* + 1) = 800 for amino acid sequences.

### Prediction Profiles

Prediction profiles were computed from feature weights as described by Palme and colleagues [30,31,37]. Prediction profiles quantify the contribution of each sequence position to the decision value (public, private). Thus, prediction profiles provide improved biological interpretability of the learning results compared to single feature weights because those individual positions or sequence stretches that drive classification accuracy most become visible [30].

### Sklearn support vector machine analysis

For public vs. private discrimination based on amino acid and V, N1, D, N2, J composition, the sklearn implementation of SVM [65] for Python [54] was employed with the cost parameter set at C=10 as determined by cross-validation.

## Acknowledgments

We thank Dr. Christian Beisel, Manuel Kohler, Ina Nissen and Elodie Burcklen from the Genomics Facility Basel of ETH Zürich for their expert technical assistance with Illumina high-throughput sequencing. We thank Sepp Hochreiter (JKU Linz, Austria) for helpful discussions. This work was funded by the Swiss National Science Foundation (Project #: 31003A_143869, to STR), SystemsX.ch – AntibodyX RTD project (to STR), Swiss Vaccine Research Institute (to STR). The professorship of STR is made possible by the generous endowment of the S. Leslie Misrock Foundation.

---

**Supplementary Figure 1.**
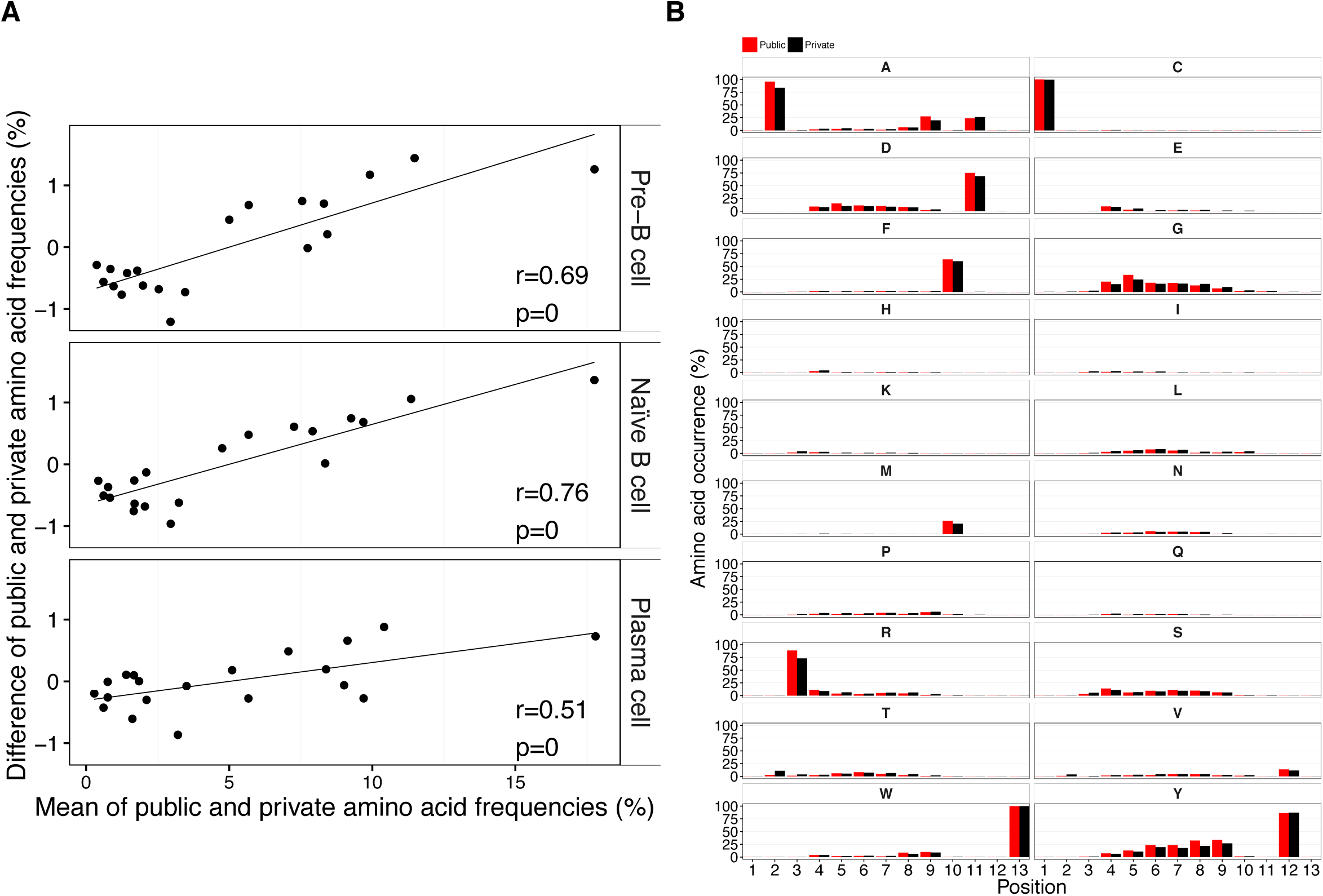
The CDR3 amino acid composition at each CDR3 sequence position differs slightly between public and private clones (Dataset 1). **(A)** Related to Figure 2D, the *difference* in public and private amino acid CDR3 frequencies is shown as a function of the *mean* of public and private CDR3 amino acid frequencies (by B-cell stage). The Pearson correlation coefficient and its p-value is shown for each B-cell stage. **(B)** The amino acid composition of public and private clones is exemplarily shown at each position for CDR3s of amino acid length 13 (preBC repertoire). Barplots show mean±s.e.m.

**Supplementary Figure 2.**
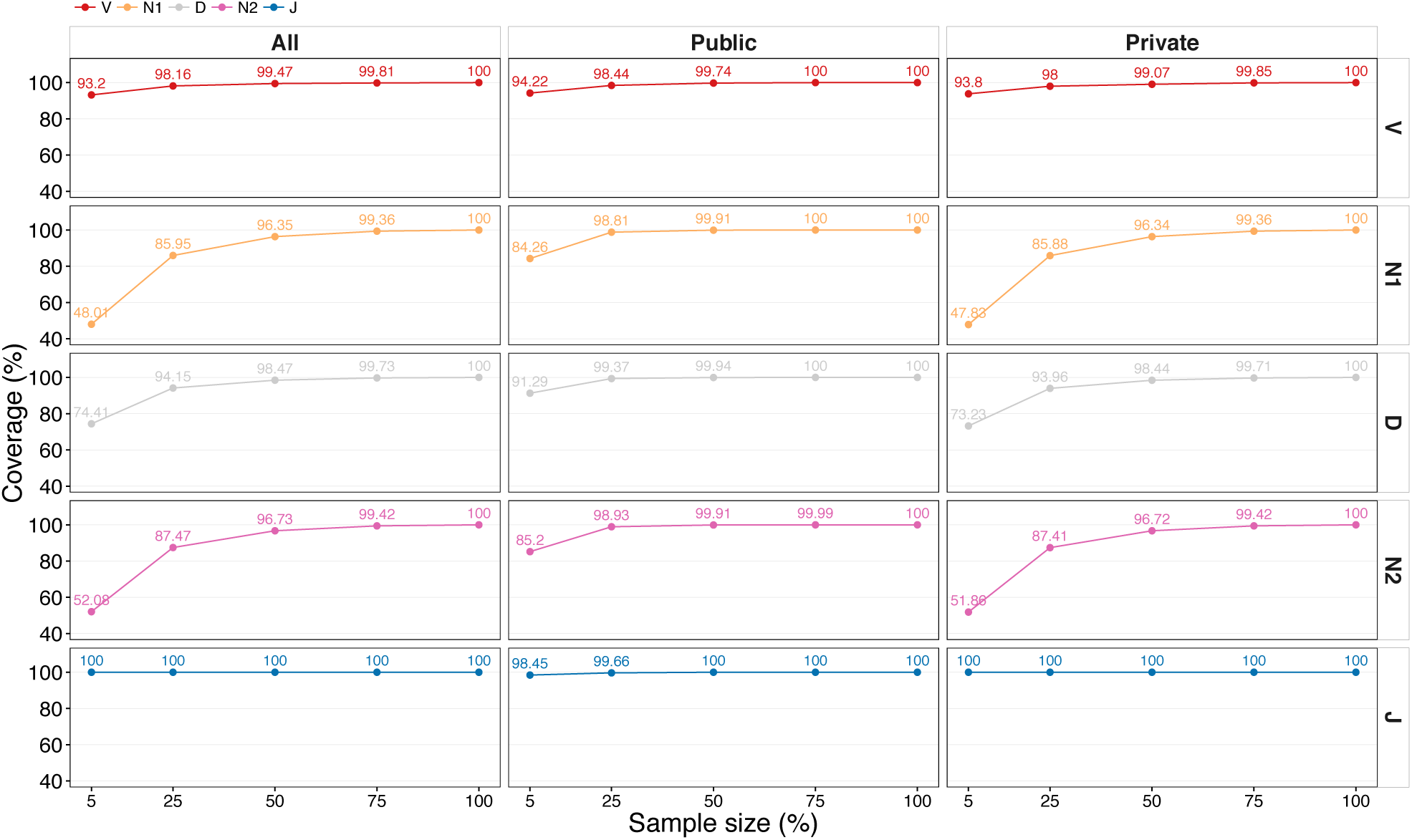
Technological coverage of each subregion is achieved at 25–50% of sequencing reads (Dataset 1). To estimate the technological coverage of each CDR3 subregion (V, N1, D, N2, J), bootstrapping was performed. 5, 25, 50, 75 and 100% of the full diversity of each subregion was sampled to subsequently compare the bootstrapped diversity to the total number of unique sequences (coverage).

**Supplementary Figure 3.**
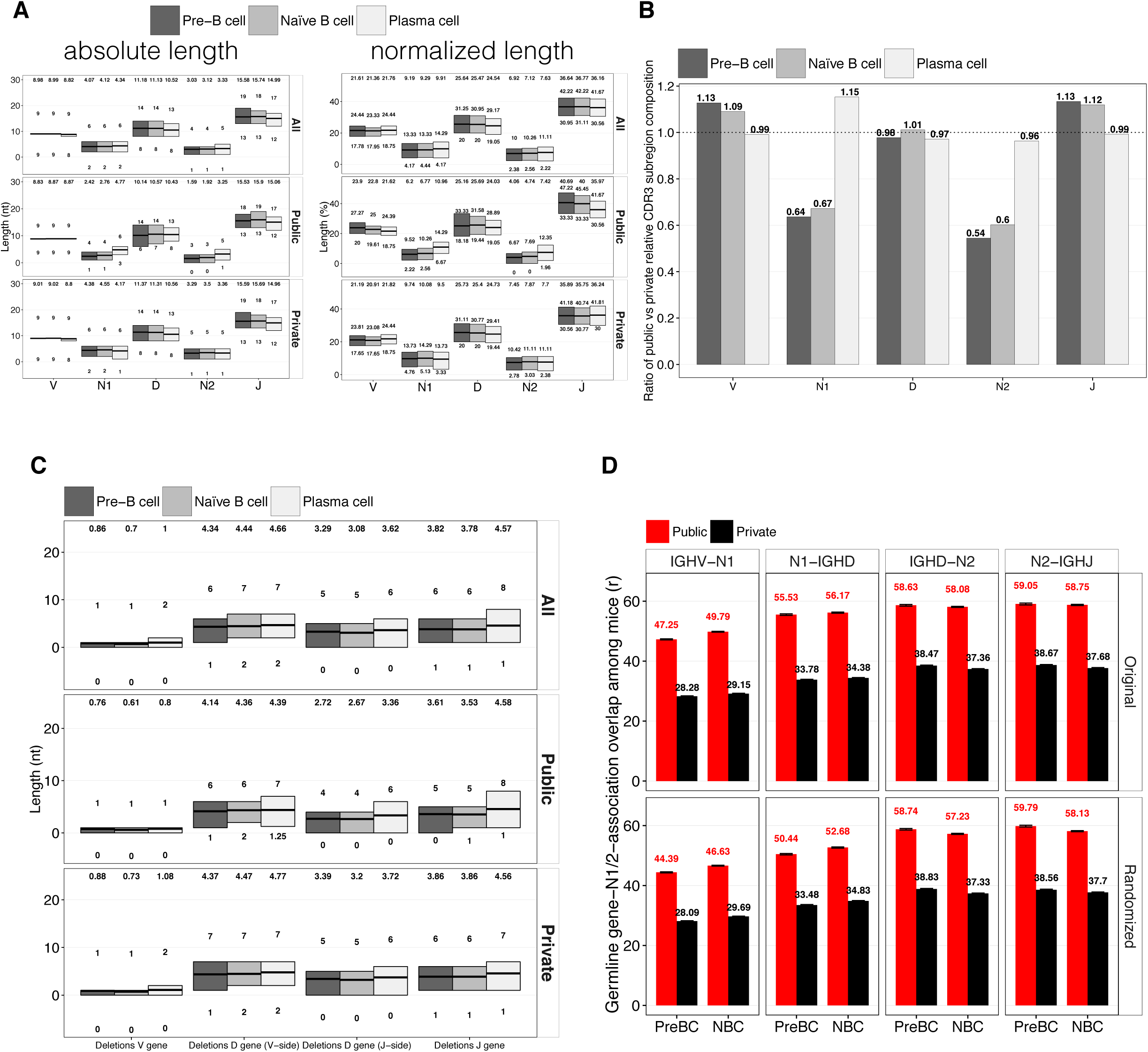
Length comparison of CDR3 subregions and deletions between public and private clones (Dataset 1). **(A)** Absolute and normalized CDR3 subregion lengths by B-cell population for public, private and all clones (irrespective of public/private status). Differences between public and private clones of preBC and nBC are significant (p<0.05). **(B)** Ratios of normalized CDR3 subregions lengths (Figure 3B). Largest deviations from 1:1 ratio between public and private clones observed in preBC for the N1 and N2 subregions (36 and 46 percentage points, respectively). **(C)** Absolute length of V, D, J deletions by B-cell population. Differences between public and private clones of preBC and nBC are significant (p<0.05). **(D)** Nucleotide insertions (N1/N2) were aggregated by germline (IGHV/D/J, panels). Subsequently, the overlap of N1/N2 insertions was compared across mice, averaged and displayed by B-cell population (original). “Randomized" means the randomization of the association of germline genes and N1/N2 insertions. Barplots show mean±s.e.m.

**Supplementary Figure 4.**
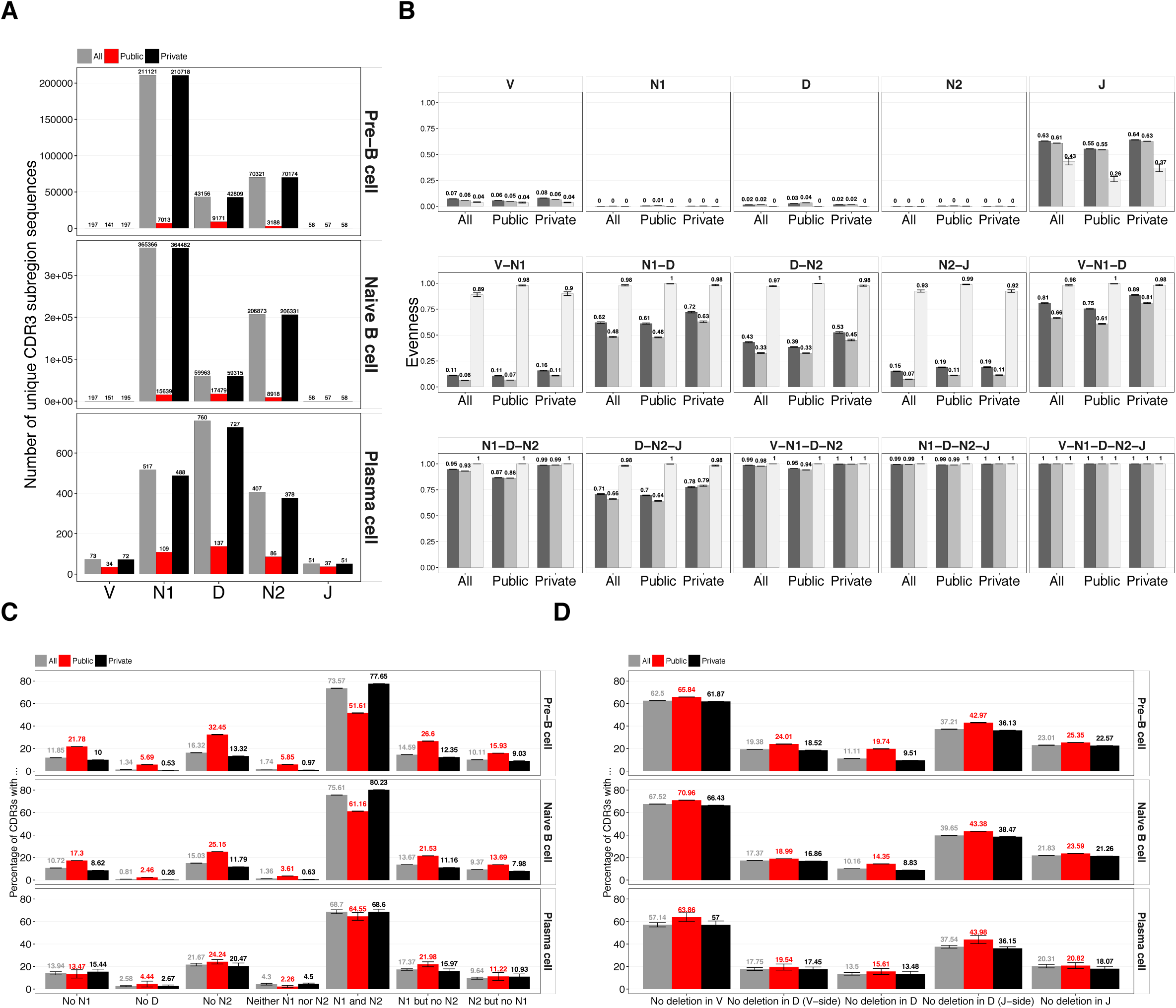
Comparison of CDR3 subregion (and deletions) diversity and occurrence between public and private clones (Dataset 1). **(A)** Total number of unique CDR3 subregion (V, N1, D, N2, J) sequences by B-cell population (non-bootstrapped, see Figure 3D for bootstrapped version). **(B)** The Shannon evenness (see Methods) of CDR3 subregions and their combinations by B-cell population. **(C)** Frequency of N1, N2 insertions across cases by B-cell population. **(D)** Frequency of deletions across cases by B-cell population. Notably, deletions consistently occurred more often in private clones with the strongest contrast found in pre-B cells (private: 90.48% (100%-9.52%) vs. public: 80.38% (100%-19.62%)). Barplots show mean±s.e.m.

**Supplementary Figure 5.**
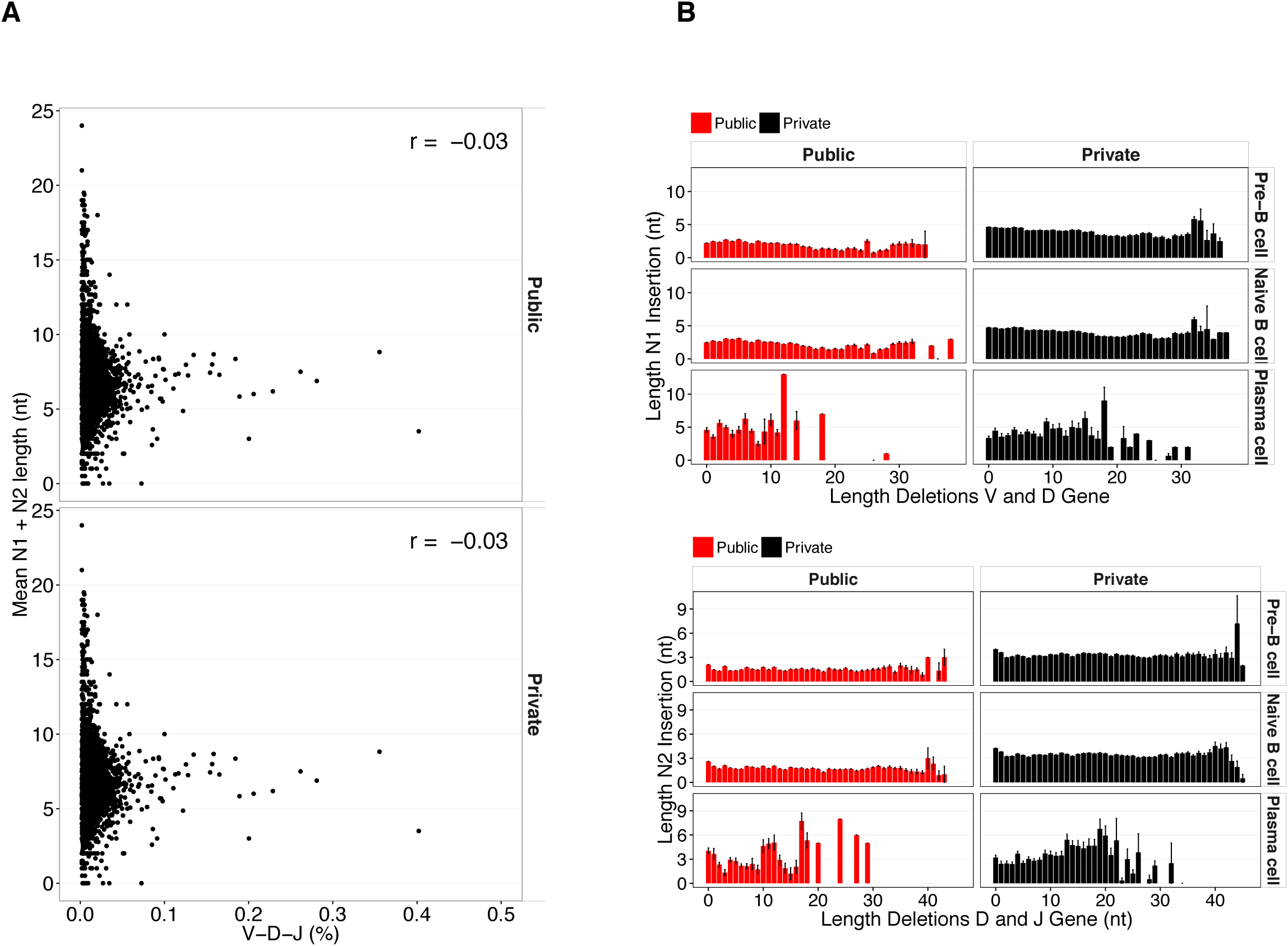
N1 and N2 subregion length is neither associated with germline gene usage nor with deletion length (Dataset 1). **(A)** The frequency of each V-D-J combination was plotted against its corresponding mean N1+N2 length. Pearson correlation is shown. **(B)** Association of N1/N2 insertions and deletions (V/D, and D/J) by B-cell population and clonal status (public/private). Barplots show mean±s.e.m.

**Supplementary Figure 6.**
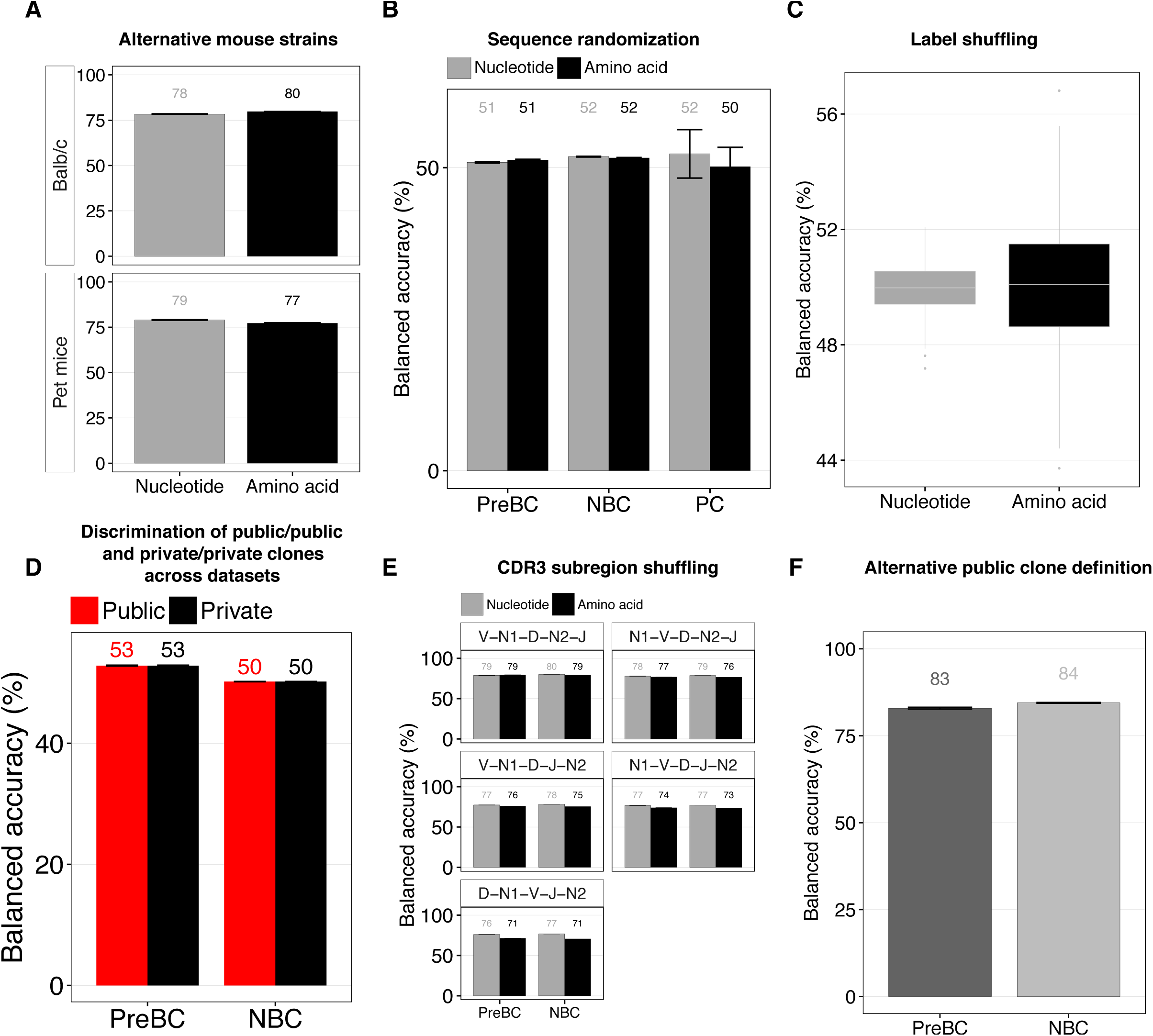
SVM-based discrimination of amino acid CDR3s and negative SVM controls. **(A)** Analogously to Figure 4A, SVM-based discrimination was performed on public and private amino acid CDR3 sequences of nBC from Balb/c (Dataset 2) and pet mice (Dataset 3). **(B)** Balanced accuracy of SVM-based discrimination of randomized public and private CDR3 sequences (nucleotide, amino acid) by B-cell population (Dataset 1). CDR3 sequences were randomized by nucleotide/amino acid shuffling. SVM was performed as described for Figure 4A. **(C)** Balanced accuracy of SVM-based discrimination performed on an nBC sample of Dataset 1 of which the labels (public, private) were shuffled in a random order. SVM was performed as described for Figure 4A. **(D)** Balanced accuracy of SVM-based discrimination of public vs. public and private vs. private clones of repertoires of identical B-cell population. SVM was performed as described for Figure 4A. This was to confirm that public and private clones across mice do not differ from one another. **(E)** Balanced accuracy of SVM-based discrimination of preBC and nBC repertoires (Dataset 1) of which the CDR3 subregions were shuffled as indicated. **(F)** Validation that public/private clone balanced prediction accuracy is independent of public clone definition. In contrast to the public clone definition adopted for Dataset 1, public clones were defined as those clones that were shared among *all* mice of a given cohort and B-cell population (size of CDR3-length equilibrated SVM datasets: 4’682 ± 657 (preBC, mean ± sd), 28’249 ± 6736 (nBC)). Subsequently, SVM-based discrimination of public and private amino acid was carried out analogously to that described for Figure 4A. Barplots show mean±s.e.m.

**Supplementary Figure 7.**
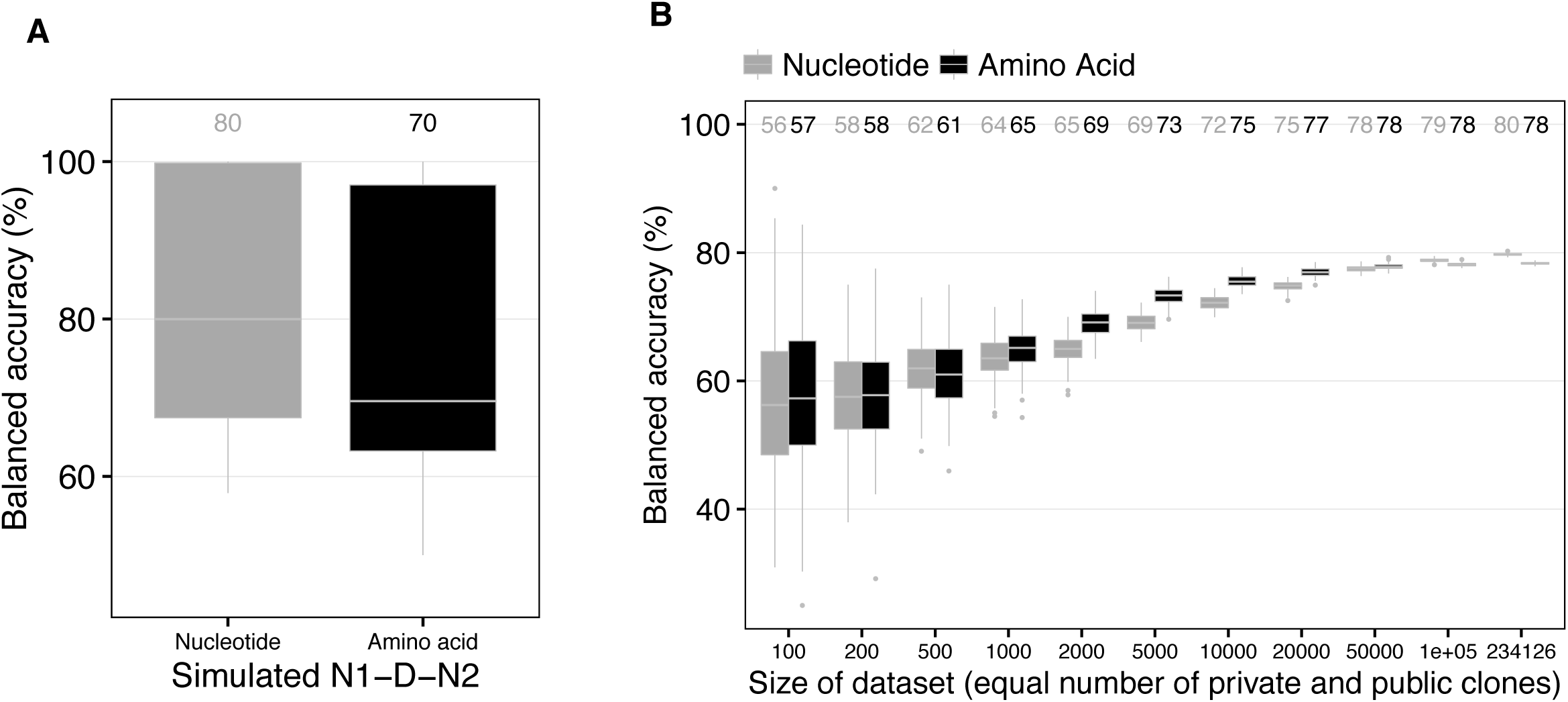
Removal of differences in N1, D, N2 subregion length between public and private clones does not influence prediction accuracy and prediction accuracy increases as a function of dataset size. **(A)** Prediction accuracy of sequence-kernel-based SVM-classification of simulated public and private N1-D-N2 sequences. In order to test whether the length differences of public and private N1, D, N2 subregions (Figure 3A) impact prediction accuracy, N1, D, N2 subregion sequences of public and private clones from one nBC repertoire (Dataset 1) were randomly assembled to N1-D-N2 simulated sequences. These simulations were performed 1’500 times: for each simulation run, the individual N1, D, N2 subregion lengths were held constant for both public and private clones and SVM parameters were chosen as for Figure 4A. **(B)** Public and private clone balanced accuracy as a function of dataset size. From the largest nBC repertoire (Dataset 1), 100–234’126 CDR3 sequences were drawn randomly 100 times to subsequently perform SVM-based prediction of public and private clones (analogously to Figure 4A). For each simulated dataset, the number of public and private clones drawn was kept identical.

**Supplementary Figure 8.**
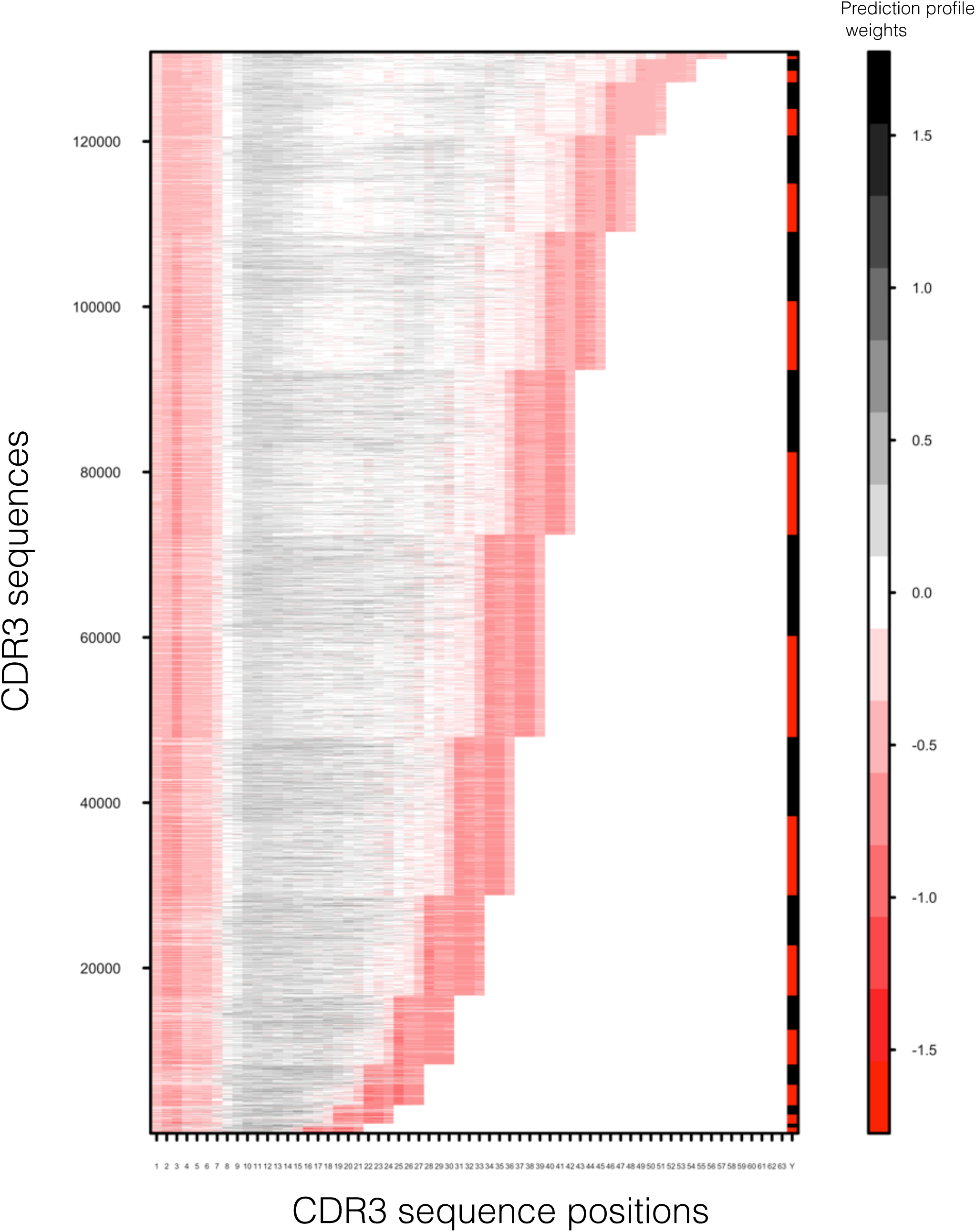
Prediction profiles are shown for all public and private CDR3 sequences of an exemplary naïve B-cell repertoire (Dataset 1). See Figure 5 and Methods for explanation of prediction profiles. Legend: red: public clone, black: private clone.

**Supplementary Figure 9.**
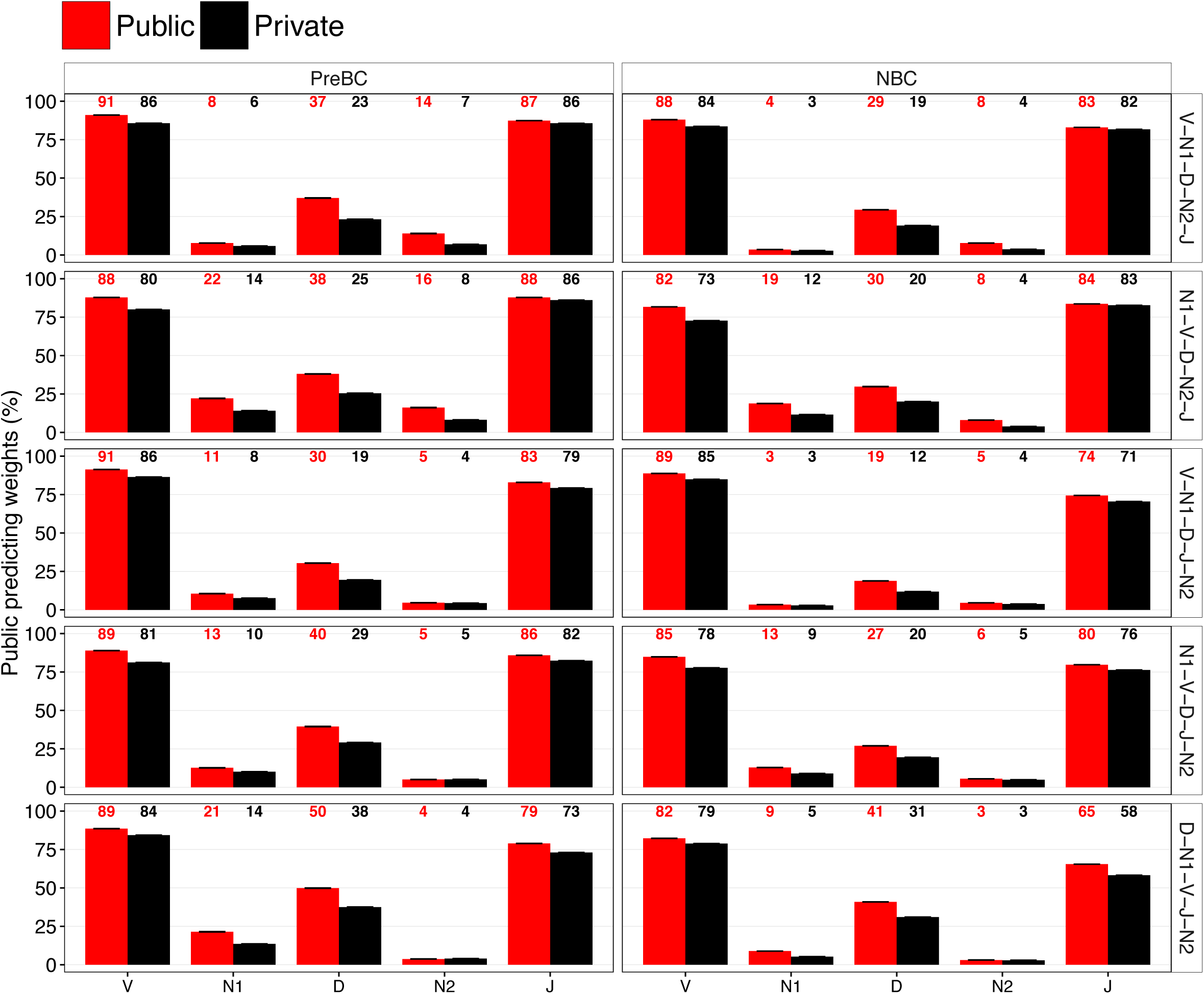
Quantitative prediction profile analysis by V, N1, D, N2, J subregion. For each of the five SVM cases shown in Supplementary Figure 6E, the percentage of public-predicting SVM weights (<0) was determined by subregion (V, N1, D, N2, J), public/private status and B-cell population across all CDR3 lengths (not solely CDR3 length 39 as shown in Figure 5B). Barplots show mean±s.e.m.

**Supplementary Figure 10.**
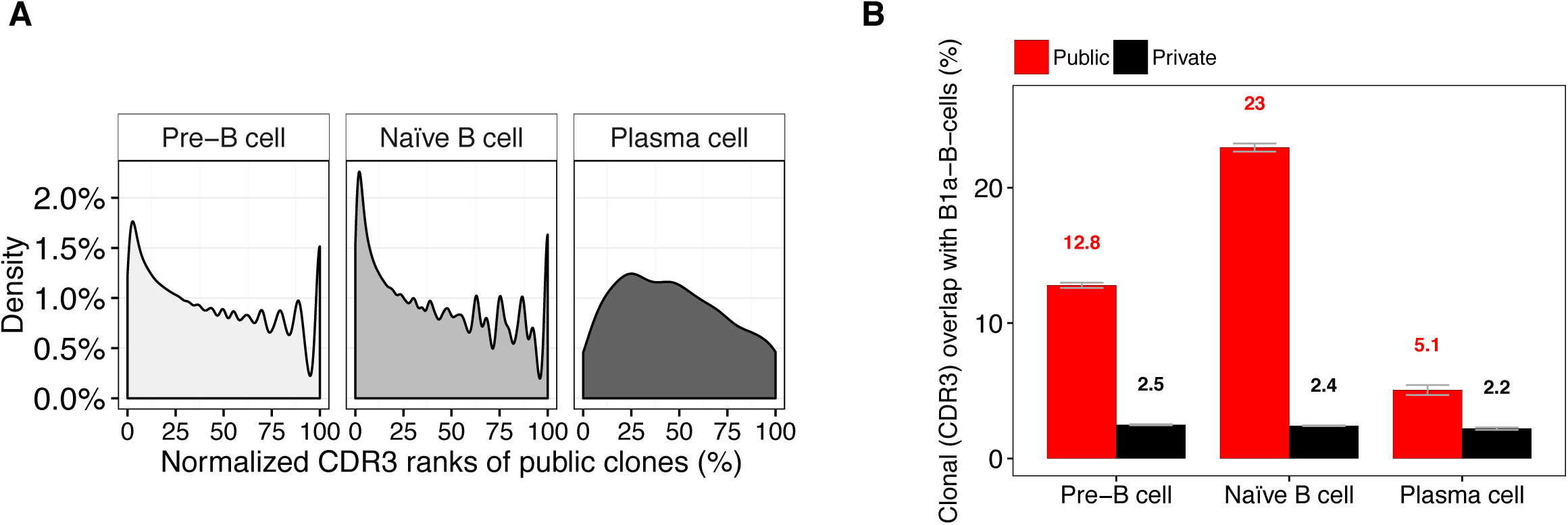
Murine B2-B-cell public clones are biased towards higher frequencies and are enriched in natural antibody producing B1a-B-cells. **(A)** Mean normalized rank of public CDR3s among all clones within a repertoire. PreBC and nBC public clones are more likely to occur at higher frequency then expected at random (uniform distribution). **(B)** PreBC, nBC and PC (Dataset 1) public and private clone overlap with B1a-B-cells (Dataset 1). Differences between public and private clones were significant (p<0.05). Barplots show mean±s.e.m.

**Supplementary Table 1.**
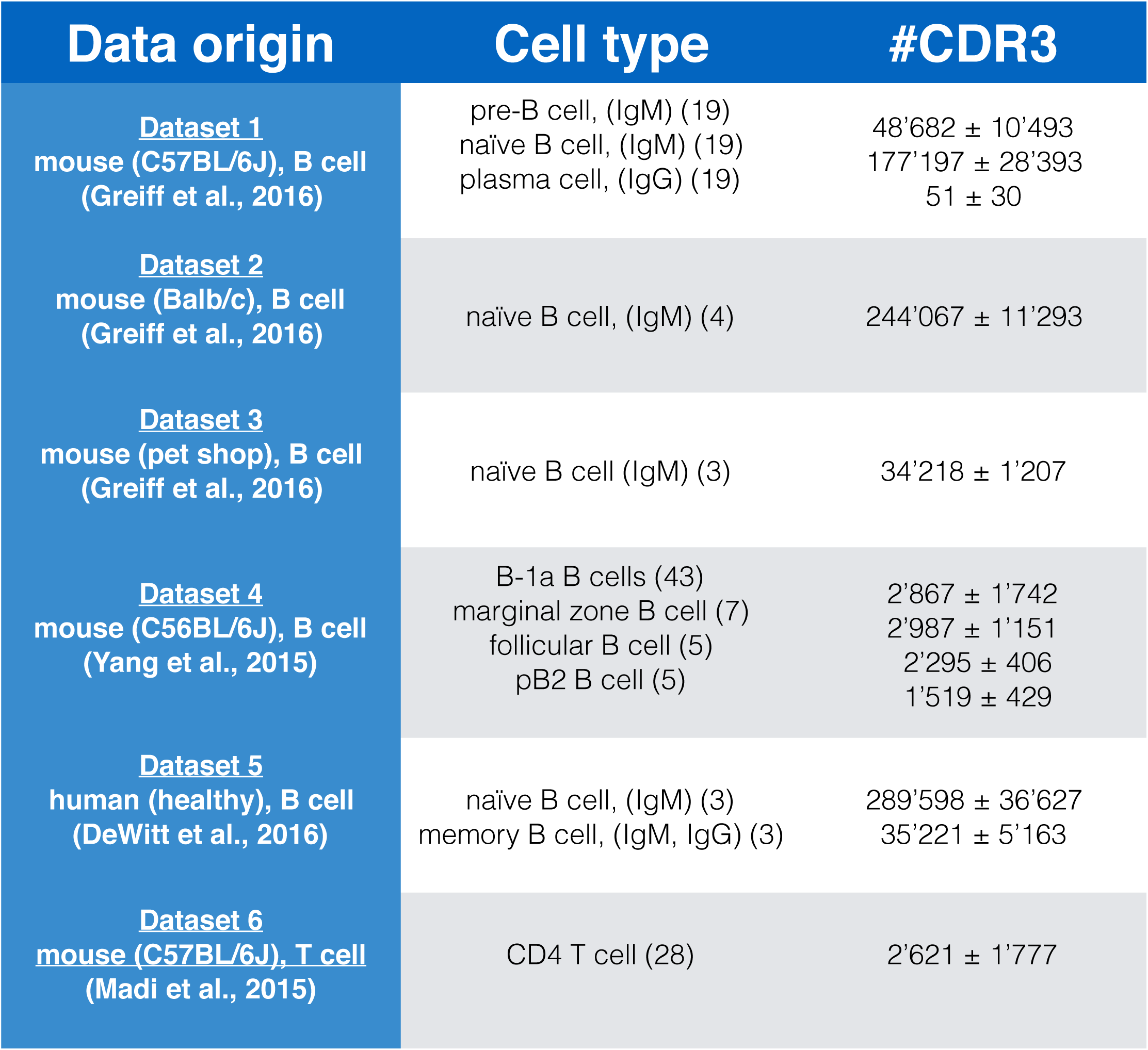
Size of CDR3-length equilibrated datasets used for SVM classification. For each of the six datasets used in this study (see Methods), a dataset of CDR3-length equilibrated sequences was constructed consisting of 50% public and 50% private CDR3 sequences. Mean and standard deviation across all samples of a given dataset and B/T cell population are displayed. Numbers in brackets indicate sample size.

| Data origin | Cell type | #CDR3 |
| --- | --- | --- |
| <u>Dataset 1</u><br>mouse (C57BL/6J), B cell<br>(Greiff et al., 2016) | pre-B cell, (IgM) (19)<br>naïve B cell, (IgM) (19)<br>plasma cell, (IgG) (19) | 48'682 ± 10'493<br>177'197 ± 28'393<br>51 ± 30 |
| <u>Dataset 2</u><br>mouse (Balb/c), B cell<br>(Greiff et al., 2016) | naïve B cell, (IgM) (4) | 244'067 ± 11'293 |
| <u>Dataset 3</u><br>mouse (pet shop), B cell<br>(Greiff et al., 2016) | naïve B cell (IgM) (3) | 34'218 ± 1'207 |
| <u>Dataset 4</u><br>mouse (C56BL/6J), B cell<br>(Yang et al., 2015) | B-1a B cells (43)<br>marginal zone B cell (7)<br>follicular B cell (5)<br>pB2 B cell (5) | 2'867 ± 1'742<br>2'987 ± 1'151<br>2'295 ± 406<br>1'519 ± 429 |
| <u>Dataset 5</u><br>human (healthy), B cell<br>(DeWitt et al., 2016) | naïve B cell, (IgM) (3)<br>memory B cell, (IgM, IgG) (3) | 289'598 ± 36'627<br>35'221 ± 5'163 |
| <u>Dataset 6</u><br>mouse (C57BL/6J), T cell<br>(Madi et al., 2015) | CD4 T cell (28) | 2'621 ± 1'777 |

**Supplementary Table 2.** CDR3 subregion diversity was sampled with >70% coverage. 1st column: cumulative (across preBC and nBC, Dataset 1) species richness of each CDR3 subregion, 2nd column: extrapolation of theoretical diversity using nonlinear regression (see Methods), 3rd column: percentage ratio of 1st and 2nd column. The approximate naïve murine nucleotide CDR3 diversity can be determined by calculating the product of the entries in the 2nd column ≈ 198 × 10^7.4^ × 10^4.7^ × 10^6.9^ × 58 ≈ 10^23^.

|  | Total number (38 preBC/<br>nBC samples) | Extrapolated<br>theoretical diversity | Coverage of extrapolated<br>theoretical diversity (%) |
| --- | --- | --- | --- |
| V | 198 | 198 | 100 |
| N1 | $10^{5.4}$ | $10^{7.4}$ | 71 |
| D | $10^{4.7}$ | $10^{4.7}$ | 100 |
| N2 | $10^{5.2}$ | $10^{6.9}$ | 74 |
| J | 58 | 58 | 100 |

